# Modulation of lysosomal Cl^-^ mediates migration and apoptosis through the TRPML1 as a lysosomal Cl^-^ sensor

**DOI:** 10.1101/2022.12.23.521794

**Authors:** Dongun Lee, Dong Min Shin, Jeong Hee Hong

## Abstract

Lysosomes are responsible for protein degradation and clearance in cellular recycling centers. It has been known that the lysosomal chloride level is enriched and involved in intrinsic lysosomal function. However, the mechanism by which chloride levels can be sensed and the chloride-mediated lysosomal function is unknown. In this study, we verified that reduced chloride levels acutely induced lysosomal calcium release through TRPML1 and lysosomal repositioning toward juxtanuclear region. Functionally, low chloride-induced lysosomal calcium release attenuated cellular migration. In addition, spontaneous exposure to low chloride levels dysregulated lysosomal biogenesis and subsequently induced the delayed migration and promoted apoptosis. Two chloride-sensing GXXXP motifs in the TRPML1 were identified. Mutations in the GXXXP motif of TRPML1 did not affect chloride levels and no changes in migratory ability. In this study, we demonstrated that the depleted chloride approach induces reformation of the lysosomal calcium pool, and subsequent dysregulated cancer progression will assist in improving therapeutic strategies for lysosomal accumulation-associated diseases or cancer cell apoptosis.

## Introduction

Cl^—^ is the most abundant electrolyte and is important for its homeostatic role in living organisms. Changes in intracellular Cl^—^ concentration ([Cl^—^]_i_) participate in the determination of membrane potential and modulation of intracellular pH and cellular volume through various Cl^—^ channels and transporters (Suh & Yuspa, 2005). Each cellular compartment possesses sophisticated Cl^—^-regulation of homeostatic cellular functions. Lysosomes contain 110 mM Cl^—^, which is more than two times higher than in the cytosol (Chakraborty *et al*, 2017; Sonawane *et al*, 2002). In recent years, high lysosomal Cl^—^ content is represented its critical role in lysosomal functions such as degradative function and other cellular processes (Chakraborty *et al*., 2017; Pu *et al*, 2016).

Lysosomal-mediated metabolic processes regulate cellular homeostasis and clearance responses. Abnormalities in lysosomal functions are involved in various diseases including lysosomal storage diseases and cancers (Dai *et al*, 2017; Festa *et al*, 2018; Goldsmith *et al*, 2014; Levine & Kroemer, 2008; Yin *et al*, 2019). Although high lysosomal Cl^—^ content is critical and the primitive role of Cl^—^ in lysosomes has been addressed in the regulation of lysosomal pH, how Cl^—^ and which protein is involved in cellular process are not well understood. Thus, we focused on the phenomena of high luminal Cl^—^ concentrations in lysosomes and depleted Cl^—^ states-mediated cellular responses. In addition, we hypothesized that the Cl^—^ modulation-based regulation of lysosomal fusion or attenuation of defective autophagosomes could be potential therapeutic strategies for lysosome-based cancer targeting.

In this study, we verified that the lysosomal Ca^2+^ channel TRPML1, activated by depleted Cl^—^, initiates autophagic flux and promotes autophagosome formation in the juxtanuclear region. We identified, for the first time, the lysosomal Cl^—^ sensing motif in TRPML1 and verified the role of Cl^—^ as a signaling ion in the lysosomal modulation. Thus, our study suggests that disturbance of Cl^—^ sensing dysregulates lysosomal process through a TRPML1-dependent mechanism.

## Results

### Acute [Ca^2+^]_i_ increase from lysosome through reduced [Cl^—^]_e_

Although [Cl^—^]_i_ is involved in homeostatic functions of cellular system, the role of Cl^—^ as a signaling ion is poorly understood. In this study, we found that stimulation with low Cl^—^ concentrations enhanced Ca^2+^ signaling. In the resting state with physiological salt solution (Reg), H1975 cells showed no signaling for intracellular Ca^2+^ concentration ([Ca^2+^]_i_) changes (Fig 1A). Decreasing Cl^—^ concentration in extracellular media by using 0 mM Cl^—^ solution induced Ca^2+^ oscillation for a short period of time (approximately 5 min) in H1975 cells (Fig 1B, C), suggesting that Ca^2+^ oscillation occurred in intracellular compartments. Exposure of decreased [Cl^—^]_e_, ranging from 0 to 20 mM, induced Ca^2+^ signaling in H1975 cells (Fig 1D). The 0 mM Cl^—^ solution did not exist physiologically. Thus, the following studies were performed with 5 mM Cl^—^ as the low Cl^—^ condition. As shown in Fig 1E, [Cl^—^]_i_ decreased before starting to reveal the [Ca^2+^]_i_ oscillation in low Cl^—^ extracellular media. In addition, the intracellular pH (pH_i_) increased before increasing [Cl^—^]_i_ and [Ca^2+^]_i_, and then gradually decreased. The cellular volume with calcein-AM fluorescence-mediated measurement did not change in H1975 cells in low Cl^—^ extracellular media (Fig 1E). To demonstrate the source of low Cl^—^-mediated [Ca^2+^]_i_ oscillations, we hypothesized that intracellular compartment lysosomes, which possess ion channels such as CLCs, two pore channel (TPC), and TRPMLs, transfer Cl^—^ and Ca^2+^ (Lee & Hong, 2020; Yin *et al*., 2019). To deplete lysosomal Ca^2+^, cells were treated with lysosomal Ca^2+^ exhausting agents, V-ATPase inhibitor bafilomycin-A1 (Baf) (Lopez-Sanjurjo *et al*, 2013) or selective lysosomal depletion agents through lysosomal membrane rupture glycyl-Phenylalanine β-naphthylamide (GPN) (Jadot *et al*, 1984). Low Cl^—^-mediated Ca^2+^ signaling did not increase after treatment with Baf or GPN; however, Ca^2+^ ionophore ionomycin-evoked Ca^2+^ signaling was observed (Fig 1F). Baf/GPN-induced Ca^2+^ signaling induced the exhaustion of lysosomal Ca^2+^; therefore, no changes in [Ca^2+^]_i_ were observed after GPN/Baf stimulation (Fig 1G, H). In addition, treatment with U18666A, which depletes lysosomal Ca^2+^ without affecting V-ATPase (Lloyd-Evans *et al*, 2008), decreased low Cl^—^-mediated Ca^2+^ signaling (Fig EV1A, EV1B). Ca^2+^ chelation with 1,2-bis(o-aminophenoxy)ethane-N,N,N′,N′-tetraacetic acid (BAPTA)-AM and Cl^—^ channel blocker with 5-Nitro-2-(3-phenylpropylamino) benzoic acid (NPPB) attenuated [Ca^2+^]_i_ and [Cl^—^]_i_ signaling in H1975 cells (Fig 1I, J). In addition, treatment with cyclopiazonic acid (CPA) or histamine, which induces the Ca^2+^ release of endoplasmic reticulum as another source of Ca^2+^ store has no effect on low Cl^—^-mediated Ca^2+^ signaling (Fig EV1C-EV1F). These results demonstrate that the low Cl^—^-mediated [Ca^2+^]_i_ oscillation was mediated from the lysosomal Ca^2+^ source.

**Figure 1.**
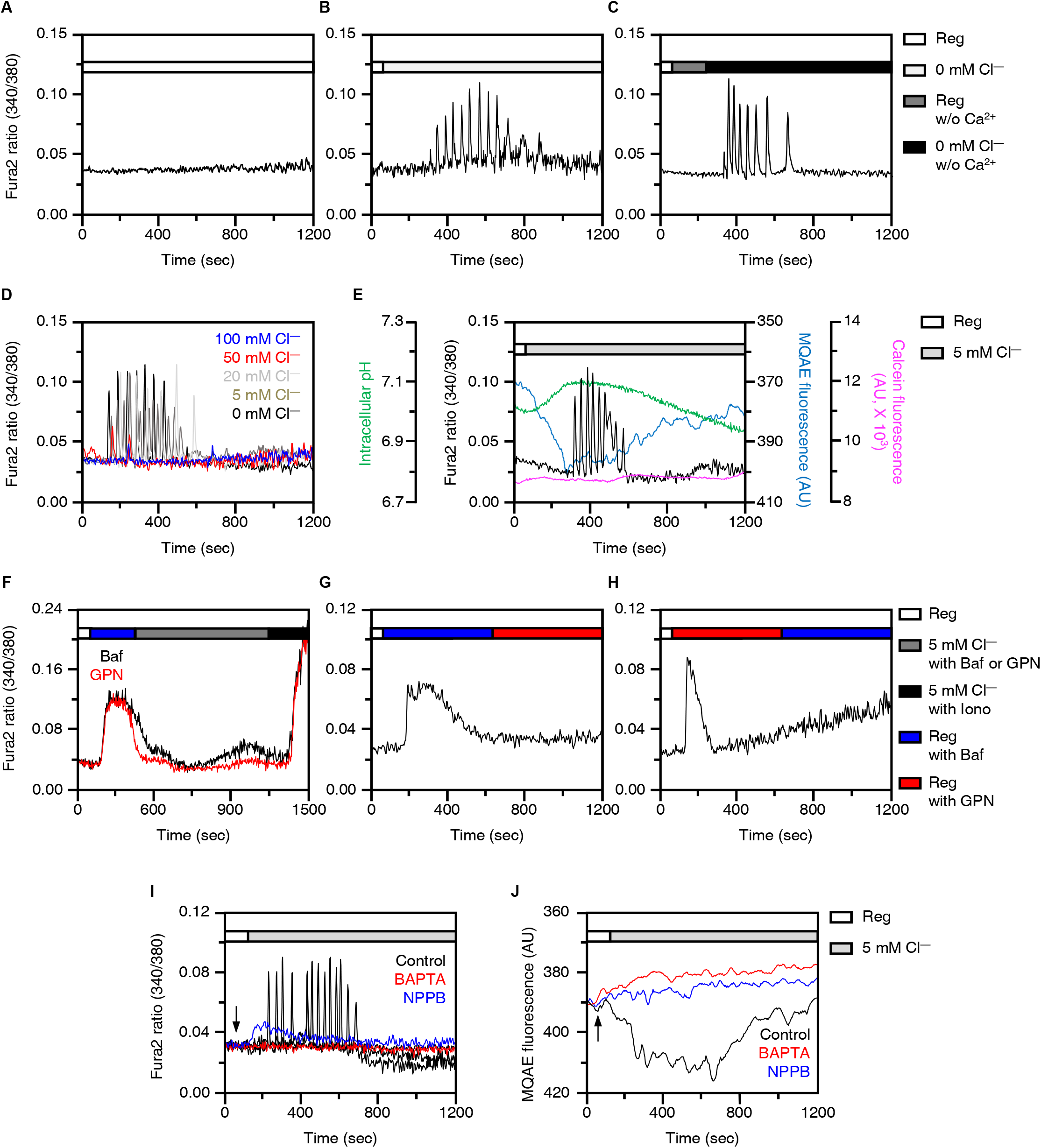
Acute [Ca^2+^]_i_ increase from lysosome through reduced [Cl^—^]_e_. **(A-D)** Changes in [Ca^2+^]_i_ in H1975 cells with Reg (A), 0 mM Cl^—^ solution (B), 0 mM Cl^—^ (Ca^2+^ free) solution (C), and solutions with various range of Cl^—^ concentration from 0 to 100 mM (D). **(E)** Combined traces of intracellular pH, Ca^2+^ (Fura2), Cl^—^ (MQAE), and cellular volume (calcein) in presence of 5 mM Cl^—^. Changes in [Ca^2+^]_i_ in H1975 cells with low Cl^—^ (5Cl^—^) in the presence of Baf (2 μM, black) or GPN (50 μM, red) which followed by treatment of ionomycin (50 μM). **(G, H)** Changes in [Ca^2+^]_i_ in H1975 cells induced by 2 μM Baf and sequential treatment of 50 μM GPN (G). Changes in [Ca^2+^]_i_ in H1975 cells induced by 50 μM GPN and sequential treatment of 2 μM Baf (H). Red and blue bars represent the time course of each reagent (red; Baf and blue; GPN). **(I, J)** Changes in [Ca^2+^]_i_ (I) and [Cl^—^]_i_ (J) with 5Cl^—^ in the presence of BAPTA-AM (10 μM, red) or NPPB (50 μM, blue). Time course of each reagent stimulation is represented with arrows. Each fluorescence measurement was proceeded with indicated conditions on the bars of right side. Each fluorescence measurement was proceeded with indicated conditions on the bars of upper side.

### Low Cl^—^-induced lysosomal reposition and deterioration of migration

Lysosomal Ca^2+^ release induces lysosomal movement toward the juxtanuclear region (Li *et al*, 2016). Attenuated [Cl^—^]_e_ and subsequent [Ca^2+^]_i_ increase induced lysosomal repositioning with the lysosomal membrane protein marker LAMP-2 staining in the perinuclear region (Fig 2A). The perinuclear clustering lysosome was located in a nuclear dent and stained with the nuclear envelope marker Lamin-A/C (Fig 2B). Treatment with BAPTA-AM and NPPB blocked lysosomal repositioning in low Cl^—^-exposed H1975 cells (Fig 2C). Low Cl^—^-mediated lysosomal repositioning was dispersed into the cytosol through re-incubation with cell culture media, called recovery (Fig 2D). These results demonstrated that low Cl^—^-mediated [Ca^2+^]_i_ oscillation induced lysosomal repositioning toward the perinuclear region with a reversible process. The acidic pH of lysosomes induces their activation to form lysosome-autophagosome fusion (Kawai *et al*, 2007; Perera & Zoncu, 2016). Transmission electron microscopy (TEM) images confirmed the low Cl^—^-induced movement of cellular vesicles toward the perinuclear region (Fig 2E). Low Cl^—^-exposed lysosomal vesicles were found in the juxtanucler region. Incubation of low Cl^—^ acidified lysosomal vesicles by enhancing pHRodo fluorescence and lysosomal acidification was inhibited by treatment with BAPTA-AM (Fig 2F, G). Lysosomal Ca^2+^ release and an acidic extracellular environment provide favorable intrinsic and extrinsic circumstances for the migration of cancer cells (Bretou *et al*, 2017; Hwang *et al*, 2019; Wu *et al*, 2021). Cancer cells are exposed to an acidic environment, and their migration of cancer cells increases in acidic pH_i_ (Hwang *et al*., 2019). The migration assay with both 4’,6-diamidino-2-phenylindole (DAPI) and crystal violet showed that H1975 cell migration was enhanced by acidic stimulation from the lower plate, whereas [Cl^—^]_e_ depletion surrounding H1975 decreased H1975 cell migration in acidic stimulation in the lower plate (Fig 2H-J). Lysosomal Ca^2+^ release induced lysosomal movement, and subsequently, lysosomal Ca^2+^ by stimulation of low Cl^—^ attenuated cellular migration.

**Figure 2.**
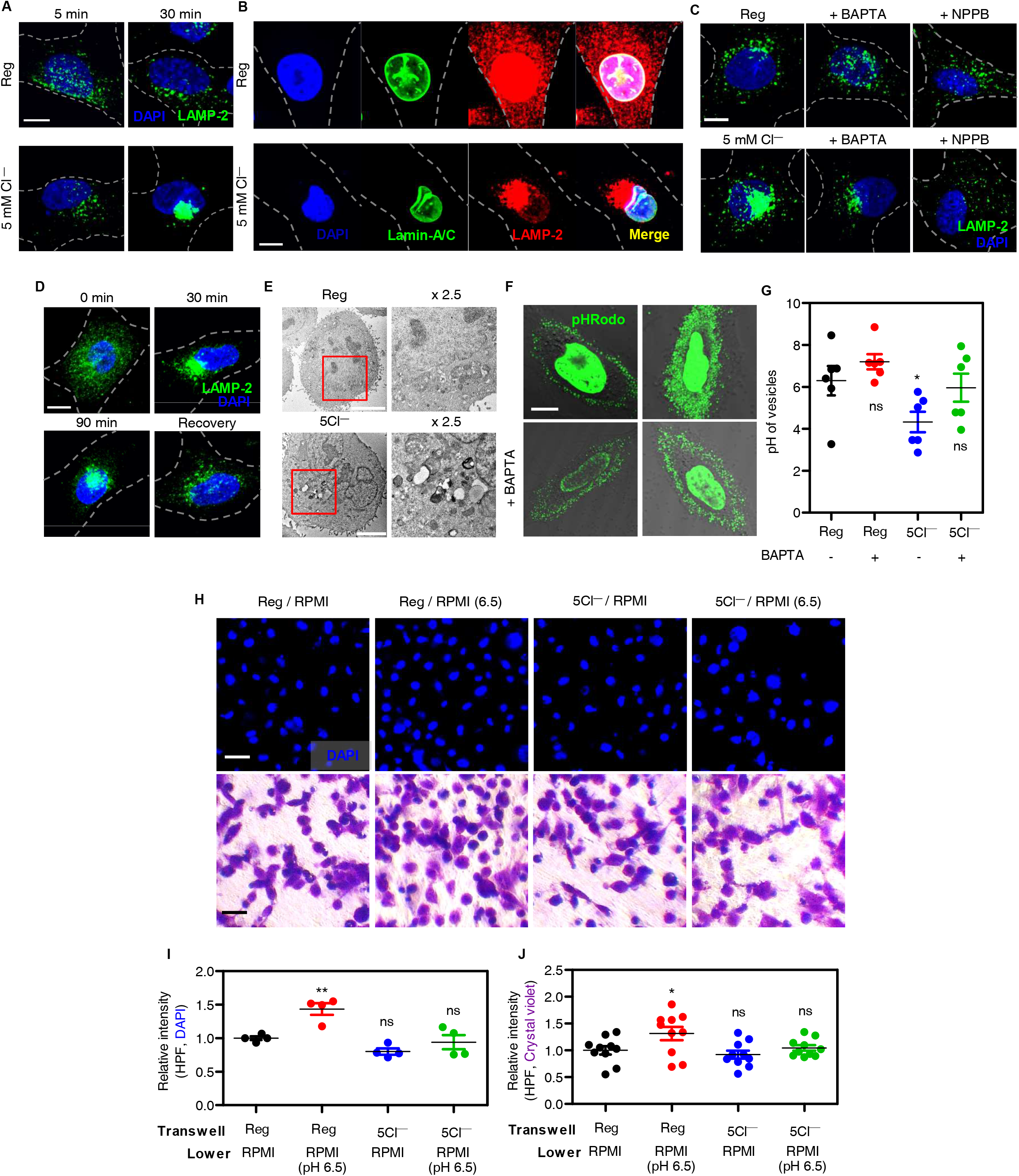
Low Cl^—^-induced lysosomal reposition and deterioration of migration. Images of immunofluorescence staining of LAMP-2 protein (green) and DAPI (blue) in H1975 cells which incubated with Reg and 5Cl^—^ solutions for 5 min and 30 min. The scale (white) represents 10 μm. **(B)** Confocal images stained for nuclear envelop (Lamin-A/C, green), LAMP-2 protein (red), and DAPI (blue) in H1975 cells which incubated with Reg and 5Cl^—^ solutions for 30 min. The scale (white) represents 10 μm. **(C)** Confocal images of the immunofluorescence staining of LAMP-2 (green) and DAPI (blue), which were incubated with Reg and 5 mM Cl^—^ (5Cl^—^) solution for 30 min in the presence or absence of BAPTA-AM and NPPB. The scale (white) is 10 μm. **(D)** Immunofluorescence images stained for LAMP-2 (green) and DAPI (blue) which incubated in indicated conditions (control; 0 min, 5Cl^—^ solution for 30 min, 5Cl^—^ solution for 90 min, and 5Cl^—^ solution for 30 min followed by incubation of media for 60 min; as called Recovery). The scale (white) represents 10 μm. **(E)** Transmission electron microscopy images for intracellular vesicles in H1975 cells with Reg or 5Cl^—^ solutions. The scale bar shows 5 μm. **(F)** Confocal microscopy images stained with pHRodo dye which incubated as indicated conditions (Reg or 5Cl^—^ in the presence or absence of BAPTA-AM). The scale (white) represents 10 μm. **(G)** The dot plots are presented as means ± SEMs of the pH value of intracellular vesicles (n=6, *p < 0.05, ns; non-significant). **(H)** Images of transwell migration assay with DAPI (blue) or crystal violet (purple) which incubated for under the indicated conditions in H1975 cells. The scale (white for DAPI and black for crystal violet) represents 50 μm. **(I)** The dot plots are presented as means ± SEMs of the relative intensity of DAPI (n=4, **p < 0.01, *p < 0.05, ns; non-significance). **(J)** The dot plots are presented as means ± SEMs of the relative intensity of crystal violet (n=10, *p < 0.05, ns; non-significance).

### TRPML1 sensed [Cl^—^]_i_ through GXXXP motif to increase lysosomal Ca^2+^ release

TRPML1-mediated Ca^2+^ release is important for lysosomal fusion in the perinuclear region (Li *et al*., 2016). Whether the changes in Cl^—^ act as a boosting signal on TRPML1 activation is still unclear. Thus, we evaluated whether TRPML1 possesses a GXXXP motif as a [Cl^—^]_i_ sensing motif, such as CLC (Dutzler *et al*, 2002; Faraldo-Gomez & Roux, 2004), Slc26a6 (Ohana *et al*, 2012), and NBCe1-B (Shcheynikov *et al*, 2015) and whether TRPML1 releases lysosomal Ca^2+^ by low Cl^—^ stimulation in the presence of a mutated GXXXP motif. Analysis of amino acid sequences showed that two GXXXP motifs were present in TRPML1. We constructed GXXXP-mutated TRPML1 mutants named as CBM (Cl^—^ binding motif)1 and CBM2 (Fig 3A, B). Lysosomal Ca^2+^ channels also include NAADP-induced TPCs (Zhang *et al*, 2011). However, the GXXXP motifs of the TPCs were analyzed in the cytosolic area (Fig EV2A). The expression of TPC1 was relatively low; therefore, we developed small interfering RNA (siRNA)-TPC2 (siTPC2-a and siTPC2-b) (Fig EV2B). Knockdown of TPC2 had no effect on low Cl^—^-mediated Ca^2+^ signaling (Fig EV2C). In addition, the GXXXP-mutated TRPM2, GP/AA mutant, had no effect on low Cl^—^-mediated Ca^2+^ signaling (Fig EV2D-F). 5 mM [Cl^—^]_e_ solution increased Ca^2+^ signaling in GP/AA-transfected H1975 cells. To verify the low Cl^—^-mediated Ca^2+^ release by TRPML1, cells were stimulated with the synthetic TRPML1 agonist ML-SA1 (Shen *et al*, 2012) after lysosomal Ca^2+^ release by low Cl^—^. ML-SA1-mediated Ca^2+^ release did not occur after low Cl^—^-mediated lysosomal Ca^2+^ depletion, and the Ca^2+^ ionophore ionomycin treatment evoked Ca^2+^ signaling after ML-SA1 (Fig 3C), suggesting that TRPML1 mediated Ca^2+^ release by treatment with low Cl^—^. Overexpressed Cl^—^-binding motif mutants, CBM1 or CBM2, revealed that low Cl^—^-mediated Ca^2+^ signaling was reduced (Fig 3D). The number of cells responding to low Cl^—^-mediated lysosomal Ca^2+^ release also decreased (Fig 3E). We confirmed the protein expression of TRPML1 mutants (Fig EV3A) and the localization of these mutants was in lysosome as co-localized with LAMP-2 (Fig EV3B-D). To confirm the channel function of the TRPML1 mutants, ML-SA1 was stimulated in the two mutants. Both CBM1 and CBM2 released Ca^2+^ upon treatment with ML-SA1, whereas in 5 mM [Cl^—^]_e_ solution, low Cl^—^-mediated Ca^2+^ signaling was reduced even in the presence of ML-SA1 (Fig 3F, G). To verify the lysosomal Ca^2+^ content in CBM1 and CBM2, we measured the Ca^2+^ increase in the presence of the selective lysosomal depletion agent GPN. GPN-induced Ca^2+^ release occurred in all TRPML1-transfected cells (Fig 3H). Although the GPN-induced Ca^2+^ release for the two mutants was slightly higher than that for the wild type, the lysosomal Ca^2+^ content was not affected. The lysosomal pH of all TRPML1 constructs, wild-type and mutant, did not change (Fig EV3E-H). To confirm whether the low Cl^—^-mediated Ca^2+^ signaling is mediated by TRPML1, we used siRNA for TRPML1, which decreases ML-SA1-induced Ca^2+^ signaling (Fig 3I, J), and the low Cl^—^-mediated Ca^2+^ signaling was decreased by siTRPML1 (Fig 3K). Treatment with ML-SA1 released lysosomal Ca^2+^, suggesting that ML-SA1-mediated lysosomal Ca^2+^ release depleted lysosomal Ca^2+^ (Fig 3L). These results indicated that the GXXXP motif is essential for the Cl^—^ sensing activity of TRPML1.

**Figure 3.**
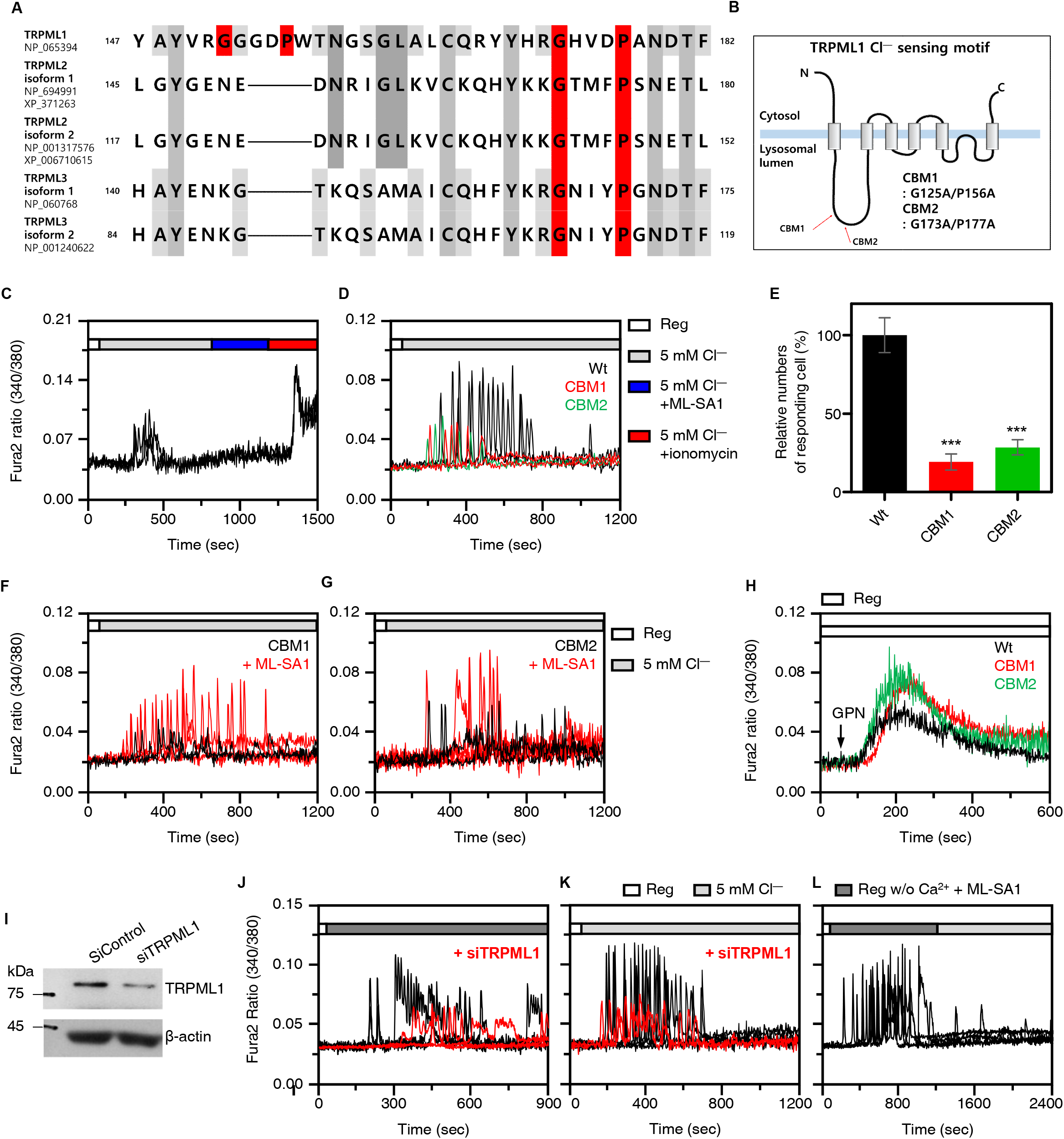
TRPML1 sensed [Cl^—^]_i_ through GXXXP motif to increase lysosomal Ca^2+^ release. Amino acid sequences of TRPML1, TRPML2, and TRPML3 isoforms marked with predicted GXXXP Cl^—^ sensing motif (red), and **(B)** schematic illustration of TRPML1 structure marked with mutation sites of GXXXP motif (CBM1 and CBM2). Changes in [Ca^2+^]_i_ in H1975 cells with 5Cl^—^ stimulation and sequential treatment of ML-SA1 (50 μM) and ionomycin (10 μM). **(D)** Changes in [Ca^2+^]_i_ with 5Cl^—^ stimulation which transfected TRPML1 wild type (Wt, black), CBM1 (red), and CBM2 (blue). **(E)** Percentage of relative numbers of responding cells as determined using R340/380 fluorescence ratios and was represented as means ± SEMs (n=3, ***p < 0.001). **(F-G)** Changes in [Ca^2+^]_i_ with 5Cl^—^ stimulation in the presence (red) or absence (black) of ML-SA1 (50 μM) which transfected CBM1 (F) and CBM2 (G). **(H)** Changes in [Ca^2+^]_i_ in H1975 cells with 5Cl^—^ stimulation and treatment of GPN which transfected TRPML1 wild type (Wt, black), CBM1 (red), and CBM2 (green). **(I)** Western blotting assay for TRPML1 in siTRPML1-transfected H1975 cells. **(J-L)** Changes in [Ca^2+^]_i_ in H1975 cells with 5Cl^—^ stimulation (J), knock down of TRPML1 (J), treatment of ML-SA1 before 5Cl^—^ stimulation (K) and treatment of 10 μM ML-SI1 (L). Each fluorescence measurement was proceeded with indicated conditions on the bars of upper side.

### Disturbed GXXXP motif of TRPML1 did not reduce cellular migration in low Cl^—^ stimulation

We focused on the reduced ML-SA1-mediated Ca^2+^ release in TRPML1 mutants. Confocal images of TRPML1 mutant-transfected H1975 cells showed enlargement of lysosomes, compared to TRPML1 wild-type transfected cells (Fig 4A, B). The fluorescence intensity profiles also showed enlargement of the lysosomes in the two mutants (Fig 4C). Transcription factor EB (TFEB) is a master regulator that translocates to the nucleus and is involved in autophagic flux and lysosomal biogenesis (Medina *et al*, 2015). In the amino acid starvation state, TFEB translocates to the nucleus and coordinates lysosomal and autophagic gene expression (Settembre *et al*, 2012). We compared the changes in TFEB expression in culture media, Reg, and low Cl^—^ solution. Enhanced TFEB translocation into the nucleus was observed at low Cl^—^ concentrations (Fig 4D, E). The CBM1- and CBM2-transfected cells showed less translocation of TFEB as compared to that in TRPML1 wild-type transfected cells (Fig 4F, G). These results indicated that the disturbed Cl^—^ sensing motif of TRPML1 mediates lysosomal dysfunction, such as enlarged lysosomes and reduced TFEB. We examined whether the disturbed Cl^—^ sensing ability of TRPML1 affected cellular migration. Cellular migration in the presence of acidic pH_e_ and treatment with ML-SA1 was enhanced. No additive effect on cellular migration was observed in either ML-SA1 or acidic conditions (Fig EV4A, EV4B). The TRPML1 mutants CBM1 and CBM2 showed reduced migration in Reg, whereas there were no changes in migration in low Cl^—^ solutions (Fig 4H-J, Fig EV4C, 4D). These results indicate that the mutated GXXXP motif disturbs the Cl^—^sensing ability of TRPML1 and reduces the migratory ability of cells.

**Figure 4.**
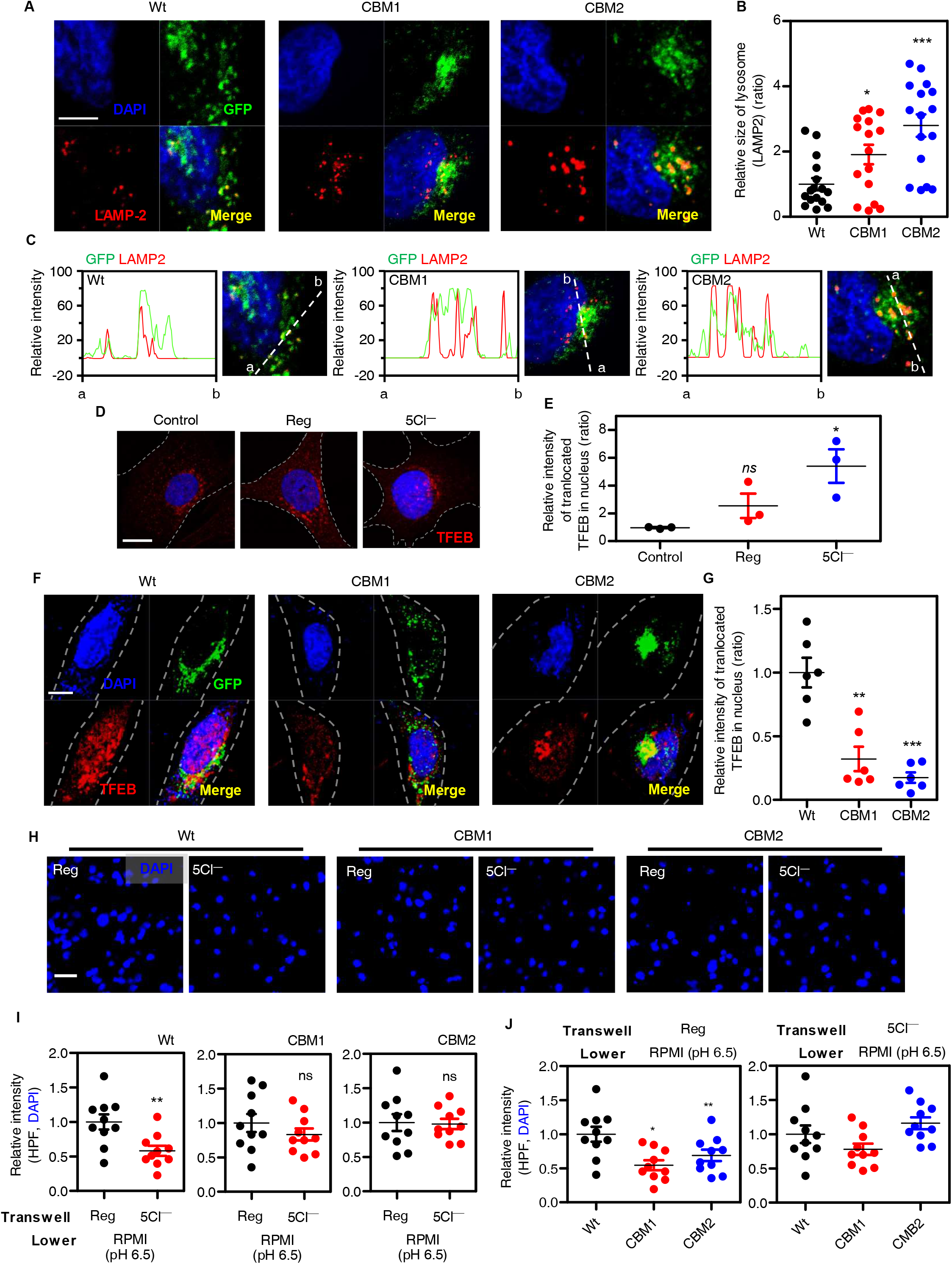
Disturbed GXXXP motif of TRPML1 did not reduce cellular migration in low Cl^—^ stimulation. Confocal images of LAMP-2 (red) and DAPI (blue) with transfection of GFP-tagged TRPML1 Wt, CBM1, and CBM2 (green). The scale (white) represents 5 μm. The dot plots are presented as means ± SEMs of the relative area of LAMP-2 (n=15, *p < 0.05, ***p < 0.001). **(C)** The fluorescence intensity profiles represented enlargement of lysosomes in GFP (green) and Rhodamine (red) signals of the indicated regions of panel A. The dotted line (white) represents region of intensity profiles between point ‘a’ and point ‘b’. **(D)** Confocal images of TFEB (red) and DAPI (blue) in 5Cl^—^ stimulation. The scale (white) represents 10 μm. **(E)** The dotted plots are presented as means ± SEMs of the relative area of LAMP-2 (n=12, *p < 0.05, ***p < 0.001). **(F)** Confocal images of immunofluorescence staining of TFEB proteins (red) and DAPI (blue) with transfection of GFP-tagged TRPML1 Wt, CBM1, and CBM2 (green). The scale (white) represents 10 μm. **(G)** The dot plots are presented as means ± SEMs of the relative intensity of translocated TFEB in nucleus (n=6, **p < 0.01, ***p < 0.001). **(H)** Images of transwell migration assay with DAPI (blue) or crystal violet (purple) in TRPML1 Wt, CBM1, and CBM2-transfected cells with the indicated conditions. The scale (white) represents 10 μm. **(I, J)** The dot plots are presented as means ± SEMs of the relative intensity of DAPI and crystal violet (n=8, *p < 0.05, *p < 0.01, ns;non-significant). (I) The dot plots are coupled with treatment of solution. (J) The dot plots are coupled with TRPML1 clones, Wt, CBM1, and CBM2.

### CLC7 is required for lysosomal Cl^—^ path and biogenesis of lysosomal proteins

Since CLC7 is expressed in late endosomes and lysosomes and is proposed to be the lysosomal Cl^—^ channel (Kasper *et al*, 2005; Kornak *et al*, 2001; Poroca *et al*, 2017), we evaluated whether depleted lysosomal Cl^—^ channel CLC7 affects TRPML1-mediated lysosomal Ca^2+^ release. CLC7 knockdown with siRNAs was evaluated based on CLC7 protein expression (Fig 5A). Silencing CLC7 with siRNAs did not mediate lysosomal Cl^—^ release to the cytosol under low Cl^—^ stimulation (Fig 5B) and reduced low Cl^—^-mediated lysosomal Ca^2+^ release (Fig 5C, D), suggesting that CLC7 is a dominant lysosomal Cl^—^ release channel in response to low Cl^—^. Whereas overexpression of CLC7 was observed no changes compared to control in Cl^—^ and Ca^2+^ measurement (Fig 5C, D). To confirm the low Cl^—^-mediated Ca^2+^ release through the dominant involvement of CLC7, the role of CLC3, which is broadly expressed, was determined. Unlike CLC7 knockdown, siRNA-CLC3 induced low Cl^—^-mediated Ca^2+^ release (Fig 5E, F). CLC3 knockdown with siRNAs was evaluated based on CLC3 protein expression (Fig EV5A). Knockdown of CLC7 reduced TRPML1 expression (Fig 5G and Fig EV5B) and lysosomal protein mTORC1 expression (total form and two types of phosphorylated form S2448, S2481 (Yim & Mizushima, 2020)) (Fig EV5B). Knockdown of TRPML1 reduced mTORC1 expression but did not reduce CLC7 expression (Fig EV5C). To confirm the reduced TRPML1 expression in siRNA-CLC7 cells, cells were stimulated with ML-SA1 without extracellular Ca^2+^ media. CLC7 knockdown did not mediate ML-SA1-activated Ca^2+^ signaling (Fig 5H), suggesting that CLC7 is important for lysosomal regulation and lysosomal protein expression. To test this hypothesis, cells were investigated for lysosomal localization in depleted CLC7. Lysosomal localization in CLC7-depleted H1975 cells was observed in the juxtanuclear region, regardless of Reg or low Cl^—^ stimulation (Fig 5I). These results indicate that CLC7 is required for lysosomal Cl^—^ transfer and biogenesis of lysosomal proteins, such as TRPML1 and mTORC1.

**Figure 5.**
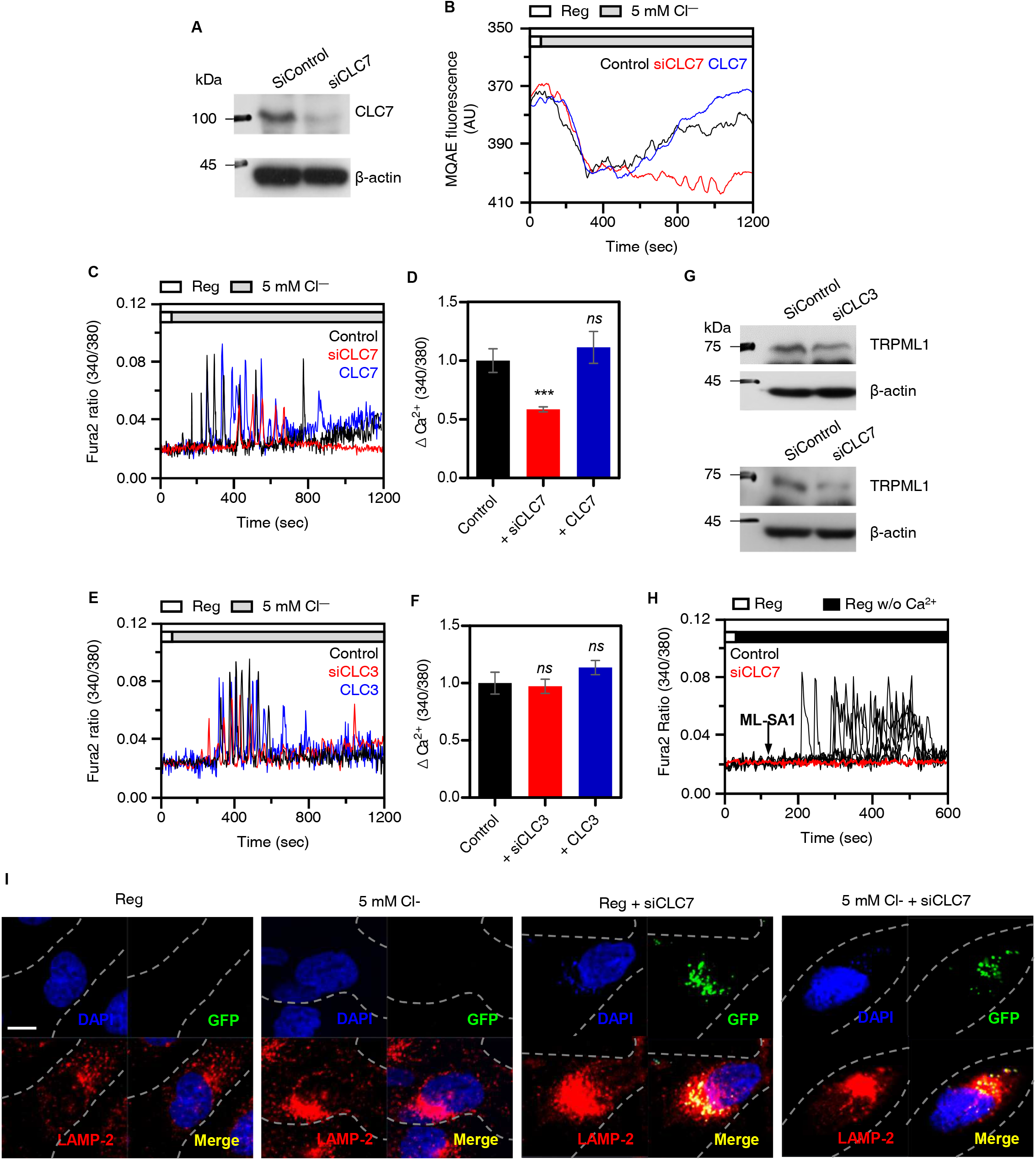
CLC7 is required for lysosomal Cl^—^ path and biogenesis of lysosomal proteins. Western blotting analysis of CLC7 transfected with siCLC7. β-actin was used as a loading control. **(B)** MQAE traces for Cl^—^ movement which silenced CLC7 (red) and over-expressed CLC7 (blue) in 5Cl^—^ stimulation **(C)** Changes in [Ca^2+^]_i_ with 5Cl^—^ stimulation which silenced CLC7 (red) and over-expressed CLC7 (blue). Analysis of maximum [Ca^2+^]_i_ peak as determined using R340/380 fluorescence ratios and means ± SEMs (n=5, ***p < 0.001, ns; non-significance). **(E)** Changes in [Ca^2+^]_i_ with low Cl^—^ stimulation which silenced CLC3 (red) and over-expressed CLC3 (blue). **(F)** Analysis of maximum [Ca^2+^]_i_ peak as determined using R340/380 fluorescence ratios and means ± SEMs (n=5, ns; non-significance). **(G)** Western blotting analysis of TRPML1 transfected with siCLC3 and siCLC7. **(H)** Changes in [Ca^2+^]_i_ in H1975 cells with 5Cl^—^ stimulation and treatment of ML-SA1 (50 μM) which transfected with siCLC7. **(I)** Immunofluorescence images stained for GFP (green), LAMP-2 (red), and DAPI (blue) which incubated as indicated conditions (Reg or 5Cl^—^ solution for 30 min) in GFP-tagged siCLC7-transfected cells. The scale (white) represents 10 μm.

### Extended low Cl^—^ treatment induces lysosomal depletion and apoptosis

As shown in fig 2H, extended treatment of low Cl^—^, which even low Cl^—^-induced lysosomal Ca^2+^ signaling is over, decreased migration of H1975 cells. The long-term application with 12 hours and 24 hours depleted lysosomes which stained with LAMP-2 in comparison with 30 min incubation (Fig 6A). To determine the effect of lysosomal depletion, apoptotic signals was determined. Cleaved PARP, a hallmark of apoptosis (Ahn *et al*, 2015; Kaufmann *et al*, 1993; Tewari *et al*, 1995), was observed in a time-dependent manner after long-term application of low Cl^—^ solution (Fig 6B, C). In addition, the expression of cleaved PARP was enhanced in Cl^—^ dose-dependent manner (0 mM to 50 mM) in accordance with lysosomal Ca^2+^ release, as described in Fig 1D (Fig 6D-E). Incubation with low Cl^—^ solution for 24 h increased late apoptosis of H1975 cells in comparison with that of the Reg-treated cells (Fig 6F, G). These results indicate that long-term treatment of low Cl^—^ induced lysosomal depletion and apoptosis.

**Figure 6.**
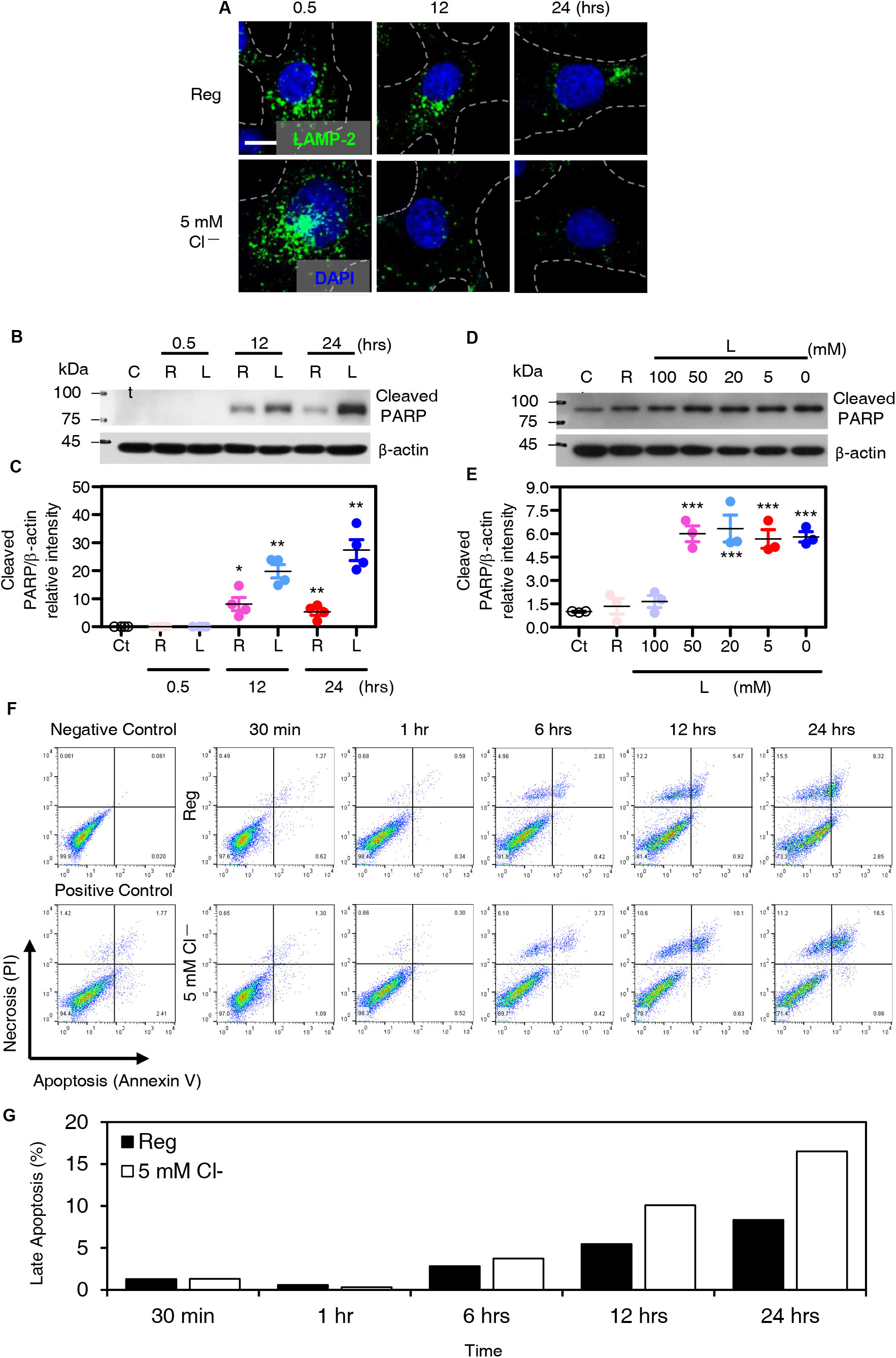
Extended low Cl^—^ treatment induces lysosomal depletion and apoptosis. Immunofluorescence images stained for LAMP-2 (green) and DAPI (blue) which incubated as indicated conditions (Reg or 5Cl^—^ solution for 30 min, 12 hrs, and 24 hrs). The scale (white) represents 10 μm. **(B)** Western blotting analysis of cleaved PARP and β-actin in H1975 cells incubated with Reg (R) and 5Cl^—^ solution (L) during long-term application of the indicated time. **(C)** The dot plots are presented as means ± SEMs of the protein band of cleaved PARP normalized to β-actin (n=4, *p < 0.05, **p < 0.01). **(D)** Western blotting analysis of cleaved PARP and β-actin in H1975 cells incubated with Reg (R) and dose-dependent low Cl^—^ solution (L, 100 mM to 0 mM) for 24 hrs. **(E)** The dot plots are presented as means ± SEMs of the protein band of cleaved PARP normalized to β-actin (n=3, ***p < 0.001). **(F)** FACS analysis of apoptosis using Pacific blue-conjugated Annexin V and PI which treated with Reg or 5 mM Cl^—^ solution at indicated time. The bars show the percentage of cells on late apoptosis.

### The similar effect of low Cl^—^ on other lung cancer cell A549

We performed most experiments on the epidermal growth factor receptor (EGFR) mutant cell line H1975. We confirmed the EGFR wild type cell lines A549 in non-small cell lung cancer. The expressions of TRPML1 mRNA and protein in H1975 cells were higher than that in another non-small cell lung cancer cell line, A549 (Fig 7A). The decreased [Cl^—^]_e_, ranging from 0 to 100 mM, induced no changes in [Ca^2+^]_i_ increase in A549 cells and less than 5 mM [Cl^—^]_e_ induced a delayed small peak of Ca^2+^ signaling compared to that of H1975 cells (Fig 7B). Lysosomal repositioning (LAMP-2 fluorescence) by low Cl^—^ stimulation was also rarely observed in A549 cells (Fig 7C). We determined whether low Cl^—^ levels mediated apoptotic signal in A549 cells. Long-term application revealed the enhanced cleaved PARP expression (Fig 7D, E). We then determined whether the relatively low lysosomal response of A549 cells compared to that of H1975 cells, induced by low Cl^—^, induces cellular migration. Migration assays using DAPI and crystal violet revealed that low Cl^—^ stimulation reduced A549 migration, as shown in H1975 cells (Fig 7F-H). Although the degree of lysosomal Ca^2+^ release differed, stimulation with low Cl^—^ also inhibited A549 migration.

**Figure 7.**
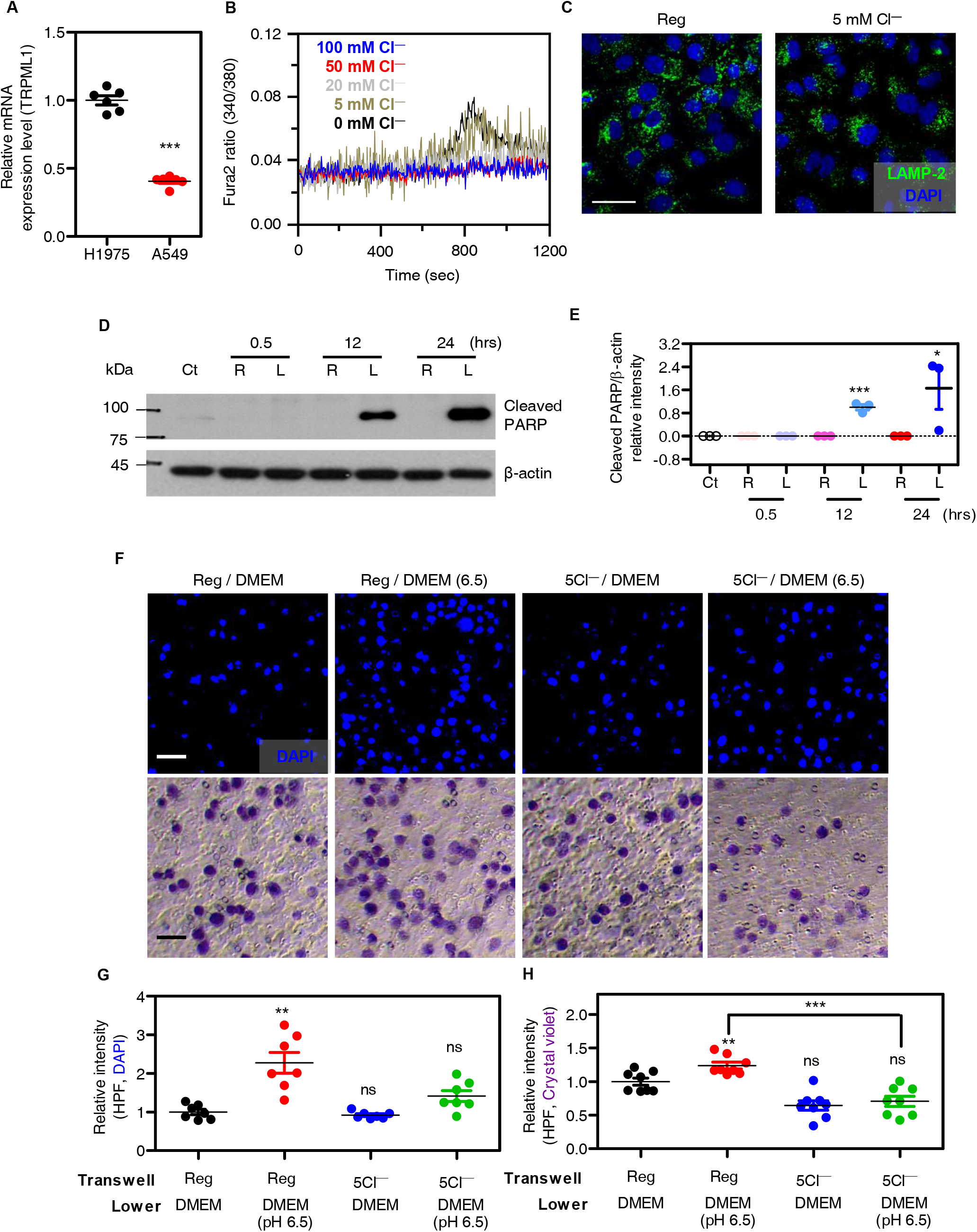
The similar effect of low Cl^—^ on other lung cancer cell A549. The dot plots are presented as means ± SEMs of TRPML1 mRNA expression in H1975 and A549 cells. **(B)** Changes in [Ca^2+^]_i_ in A549 cells with various range of Cl^—^ concentration from 0 to 100 mM. **(C)** Images of immunofluorescence staining of LAMP-2 (green) and DAPI (blue) in A549 cells which incubated with Reg and 5Cl^—^ solutions for 30 min. The scale (white) represents 25 μm. **(D)** Western blotting analysis of cleaved PARP and β-actin in A549 cells incubated with Reg (R) and 5Cl^—^ solution (L) during the indicated time. **(E)** The dot plots are presented as means ± SEMs of the protein band normalized to β-actin (n=3, *p < 0.05, ***p < 0.001). **(F)** Images of transwell migration assay with DAPI (blue) or crystal violet (purple) which incubated for under the indicated conditions in A549 cells. The scale (white for DAPI and black for crystal violet) represents 50 μm. **(G)** The dot plots are presented as means ± SEMs of the relative intensity of DAPI and crystal violet (n=4, **p < 0.01, ***p < 0.001, ns; non-significance).

## Discussion

TRPML1 is a nonselective cation channel involved in various cellular functions in migration and lysosomal biogenesis through endo/lysosomal Ca^2+^ release (Kiselyov *et al*, 2011; Medina *et al*, 2011; Yamaguchi *et al*, 2011). It is well established that TRPML1-mediated Ca^2+^ release is involved in lysosomal fusion and retrograde movement to the juxtanuclear region (Di Paola *et al*, 2018; Pu *et al*., 2016). TRPML1 modulation has been addressed in the stimulation of phosphatidylinositol (3,5) bisphosphate (PI(3,5)P2) (Dong *et al*, 2010), acidic pH (Li *et al*, 2017; Xu *et al*, 2007), reactive oxygen species (Zhang *et al*, 2016), and the OCRL gene (De Leo *et al*, 2016). Here, we addressed the changes in Cl^—^ concentration and modulated TRPML1-mediated Ca^2+^ release and TRPML1 trafficking. Depleted or reduced Cl^—^ mediated acute lysosomal Ca^2+^ release through TRPML1 and long-term exposure to reduced Cl^—^ enhanced apoptosis through lysosomal Ca^2+^ pool depletion. The two GXXXP motifs of TRPML1 in the lysosomal lumen sensed lysosomal Cl^—^ concentration, and its mutation reduced TFEB translocation and cellular migration.

Cl^—^ is a homeostatic and signaling ion (Luscher *et al*, 2020). As several features of signaling ion, reduction of Cl^—^ concentration enhanced endosome Ca^2+^ channel activity in early endosome (Saito *et al*, 2007). Our previous study addressed Cl^—^ as a modulating factor for NBCe1-B activity (Shcheynikov *et al*., 2015). Additionally, the Cl^—^ sensing role may possess in GXXXP motif-predicted various Cl^—^ transporters and channels, such as the Slc26 family, Na^+^-K^+^-Cl^—^ cotransporter, Na^+^-Cl^—^ cotransporter, and cystic fibrosis transmembrane conductance regulator (Shcheynikov *et al*., 2015). Modulation of high lysosomal Cl^—^ has been suggested to maintain lysosomal function (Chakraborty *et al*., 2017). However, whether lysosomal Cl^—^ affects TRPML1-mediated Ca^2+^ release remains relatively unknown. Reduction of Cl^—^ results in enhanced lysosomal H^+^ ions through the involvement of Ca^2+^/H^+^ exchanger, V-type ATPase, and CLC-7 (Schwartz & Muallem, 2019). Thus, low Cl^—^ induces H^+^ influx into the lysosomal lumen and subsequent Ca^2+^ release through TRPML1, likely due to the secondary effect of lysosomal acidification (Cheng *et al*, 2010). However, the reduced Ca^2+^ response by low Cl^—^ in two GXXXP mutants of TRPML1 without changes in lysosomal vesicle pH indicate that reduced Cl^—^ itself could be considered as a TRPML1 activator. Moreover, we addressed the predicted Cl^—^-sensing motif in TPCs, including TRPML1. However, TRPMLs and TPCs have different functions in endo/lysosomes (Yamaguchi *et al*., 2011). The difference in the Cl^—^ sensing position may provide diverse functions between the two types of lysosomal Ca^2+^ channels, TRPML1 and TPCs. Furthermore, in agreement with other cellular stresses, such as nutrient starvation or reactive oxygen species to activate TRPML1, low Cl^—^ can be a regulatory signaling factor of TRPML1 activity and lysosomal function.

The role of Cl^—^ as a signaling ion in lysosomes is unknown. Notably, we provided evidence that the luminal GXXXP motif of TRPML1 sensed changes in intracellular Cl^—^ acutely (less than 30 min), and subsequent TRPML1-mediated Ca^2+^ release allows lysosomal movement (Fig 8). Lysosomal repositioning is regulated by various signaling pathways, such as nutrient deficiency (Korolchuk *et al*, 2011; Starling *et al*, 2016; Willett *et al*, 2017), pathologic infection by pathogens (D’Costa *et al*, 2015; Dumont *et al*, 2010; Knodler & Steele-Mortimer, 2005), and cellular changes or stresses (Heuser, 1989; Parton *et al*, 1991; Yu *et al*, 2016; Zaarur *et al*, 2014). We observed dynamic phenomena of lysosomal movement over a short period of time. Without nutrient starvation, perhaps before the recognition of starvation, depleted Cl^—^ is sufficient to drive dynamic lysosomal trafficking with Ca^2+^ involvement. In addition, low Cl^—^-mediated acute lysosomal Ca^2+^ release was used in the juxtanuclear clustering of lysosomes, which is a hallmark of autophagy induction (Li *et al*., 2016). Acidification of lysosomes and prolonged low Cl^—^ conditions reduced migration and enhanced apoptosis. Lysosomal Ca^2+^ release is known to play a crucial role in cellular migration and progression (Yin *et al*., 2019) and the lysosomal Ca^2+^ channel TRPML1 provides a Ca^2+^ source to induce autophagic vesicles (Scotto Rosato *et al*, 2019). Although it is clear that a role TRPML1 in cell survival and expansion, interestingly, enriched lysosomal Cl^—^ levels are crucial and reduced Cl^—^ is acutely monitored by TRPML1 through the GXXXP motif. Its low Cl^—^-mediated Ca^2+^ release and lysosomal trafficking may provide a survival strategy against Cl– depletion within a short period of time.

**Figure 8.**
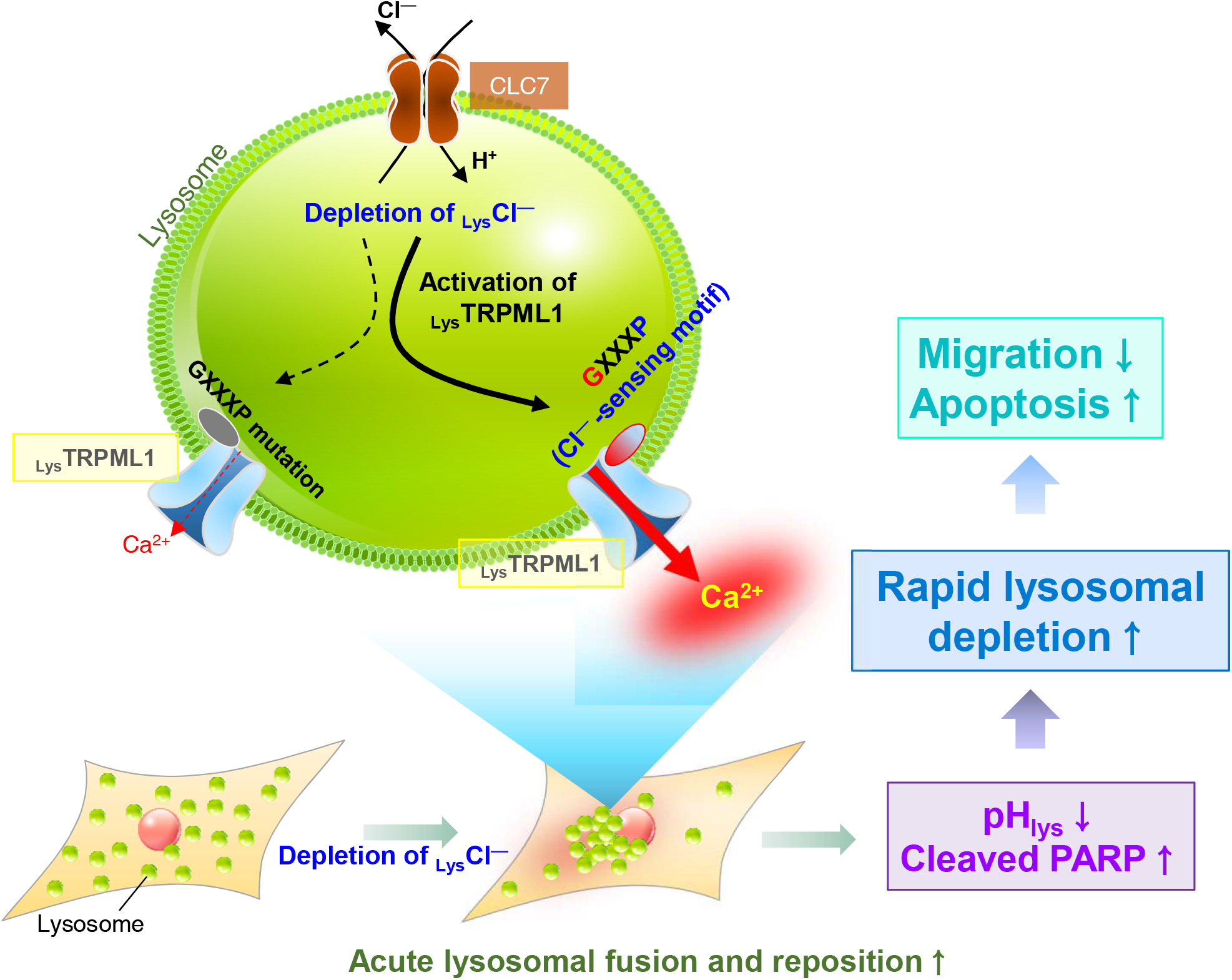
The schematic illustration of role of Cl^—^-dependent lysosomal Ca^2+^ release through TRPML1. Schematic illustration of the mechanism of Cl^—^mediated lysosomal function, including TRPML1 activation and lysosomal reposition. The GXXXP motifs of TRPML1 sense the lysosomal Cl^—^ concentration. The short period of low Cl^—^ mediated Ca^2+^ drives the juxtanuclear clustering of lysosomes. Long-term exposure to low Cl^—^ mediates excessive autophagy, decreases migration, and increases apoptosis.

Lysosomal Ca^2+^ release by TRPML1 activating signals, such as low Cl^—^ in the current study and subsequent notable changes in the lysosome-mediated mechanism, can be more potent in regulating cellular fate. The mechanism of dysregulated autophagy in several lysosomal diseases remains unclear. Lysosomal Cl^—^ has evolved from obscurity to progress in the fascinating field of autophagy. Regardless of lysosome-mediated autophagy modulation, this study attempts to explain that low Cl^—^ stimulation could be a direct and acute tool to verify lysosomal Ca^2+^ storage. On the other hand, autophagy in cancer has been considered contradictory because of its diverse effects (Rubinsztein *et al*, 2012; Yang *et al*, 2011; Yu *et al*., 2016). As we addressed in this study, low Cl^—^ approach induced depletion of the lysosomal pool and subsequent excessive autophagy through acute lysosomal Ca^2+^ release through TRPML1, dysregulated lysosomal biogenesis, and destruction of the terminal recycling center. It thus appears that Cl^—^ depletion targeting approaches, such as putative lysosomal Cl^—^ quencher, are the beginning point of improved therapeutic strategies for lysosomal targets of cancer and dysfunction of autophagy-mediated diseases.

## Material and methods

### Reagents and cell culture

Fura-2-AM (Fura2) and BCECF-AM were purchased from TEFlabs (Cat:0102 and 4011 B; Austin, TX, USA). Calcein-AM was purchased from Thermo (Waltham, MA, USA; Cat: C1430) and 1-Ethoxycarbonylmethyl-6-methoxyquinolinium bromide (MQAE) was purchased from Cayman Chemical (Ann Arbor, MI, USA; Cat:19585). Pluronic acid (Pluronic F-127, 20%(v/v) in DMSO) was purchased from Invitrogen (Carlsbad, CA, USA; Cat: P-3000MP). BAPTA-AM was purchased from Invitrogen (Cat: B1205) and CPA was purchased from Alomone Labs (Israel; Cat: C-750). Baf (from *Streptomyces griseus*), GPN, histamine, NPPB, ML-SA1, ionomycin and U18666a were purchased from Sigma (St Louis, MO, USA; Cat: B1793, G9512, D9754, H7125, N4779, SML0627, I9657, and 662015, respectively). Non-small cell lung cancer cell lines H1975 and A549 were obtained from American Type Culture Collection (Rockville, MD, USA; Cat: CRL-5908, Cat: CCL-185). H1975 cells were maintained in Roswell Park Memorial Institute 1640 (RPMI 1640, Invitrogen, Cat:11875-093) and A549 cells were maintained in Dulbecco’s modified Eagle’s medium (Invitrogen, Cat:11995-065) each containing 10% fetal bovine serum (FBS, Invitrogen; Cat:16000-044) and 100 U/mL penicillin-streptomycin (Invitrogen, Cat:15140122). Cells were incubated in a humidified incubator with 5% CO_2_ and 95% air at 37°C. When the cells reached 80% confluence, they were washed with Dulbecco’s phosphate-buffered saline (Welgene, Korea; Cat: LB001-02) after the culture medium was removed. The cells were treated with trypsin/EDTA (Invitrogen; Cat:25200-072) for 5 min. The detached cells were transferred to fresh culture dishes with coverslips for the measurement of fluorescence dye-based electrolyte changes ([Ca^2+^]_i_, [Cl^—^]_i_, pH_i_, and cellular volume) and confocal microscopy or fresh culture dishes for western blotting, flow cytometry, and migration assays.

### Plasmid, siRNA, mutation and DNA transfection

The plasmids encoding GFP-tagged Cl^—^ channels CLC3, CLC7, TRPM2, and TRPML1 were purchased from Origene (Rockville, MD, USA; Cat: RG221294, RG203450, RG216220, and RG201010). The siRNAs for CLC3 and CLC7 were constructed using double-promoter pFIV-H1/U6 siRNA cloning and expression vectors (System Biosciences, Palo Alto, CA, USA; Cat: SI111A-1), according to the manufacturer’s instructions. The vectors contained single-stranded DNA for human siRNA-CLC3 (sense, 5’-GCT GTG ATA GCC TTC CCT AAT CCA TAC-3’ and anti, 5’-TAT GGA TTA GGG AAG GCT ATC ACA GC-3’) and human siRNA-CLC7 (sense, 5’-GAT GAT CCA CTC AGG TTC AGT GAT TGC-3’ and anti, 5’-GCA ATC ACT GAA CCT GAG TGG ATC ATC-3’). The siRNAs of TRPML1, TPC1, and TPC2 were constructed by Genolution (Seoul, Korea), and the sequences were 5’-GAUCACGUUUGACAACAAA-3’ (siTRPML1), 5’-GGAGUUACCUCGUCUUUCUUU-3’ (siTPC1-a), 5’-GCUUUGUGACCCUGUUUGAUU-3’ (siTPC1-b), 5’-GGCUUUACCGACGGUAUUAUU-3’, and 5’-CCAUCAUUGGGAUCAACUUUU-3’ (siTPC2-b). The TRPM2 and TRPML1 mutant forms were produced using the QuickChange II Site-Directed Mutagenesis Kit (Agilent Technologies, Santa Clara, CA, USA; Cat:200523), according to the manufacturer’s instructions mentioned on the kit. Mutants of the Cl^—^-interacting motif GXXXP (Dutzler *et al*., 2002) were constructed to replace glycine and proline with alanine, which are indicated as GP/AA (G1390A/P1394A, for TRPM2), CBM1 (G152A/P156A, for TRPML1), and CBM2 (G173A/P177A, for TRPML1), respectively. The plasmids were transfected using jetPRIME transfection reagent (Polyplus-transfection, New York, NY, USA; Cat:114-15) according to the manufacturer’s protocol. Each plasmid was diluted in 200 μL of jetPRIME buffer, and 4 μL of the reagent was added. The mixture was incubated for 10 min at room temperature (RT) and then transferred to H1975 cells with fresh RPMI media. After 6 h of incubation, the medium was replaced with fresh medium and the cells were incubated for 24 h (CLC3, CLC7, TRPML1, CBM1, and CBM2) or 48 h (siRNA-CLC3 and siRNA-CLC7).

### Real time polymerase chain reaction (PCR)

The mRNA expression of TRPML1 in lung cancer cells (H1975, A549 and H1299) was analyzed using real-time PCR. Total RNA was extracted using GeneAll Hybrid-R™ (305-101), according to the manufacturer’s instructions. cDNA was synthesized using the AccuPower RocketScript™ Cycle RT PreMix (Bionner, K-2204). RNA (1 μg) was mixed with the premix, and the PCR cycle was 4°C (5 min) → 37°C (90 min) → 95°C (2 min). The synthesized cDNA was amplified using PowerUp™ SYBR™ Green Master Mix (Applied Biosystems, A25741). The primer sequences used were 5’-ACACCCCCAGAAGAGGAAGA-3’ (forward) / 5’-CGCAGGGACTCATGAAAAAG-3’ (reverse) for TRPML1 and 5’- GACCTGACCTGCCGTCTAGAAA-3’ (forward) / 5’-CCTGCTTCACCACCTTCTTGA-3’ (reverse) for GAPDH. The PCR cycles were as follows: 50°C (2 min) → 95°C (2 min) → [95°C (15 s) → 57°C (15 s) → 72°C (1 min)] × 40 cycles → 95°C (15 s) → 60°C (1 min) → 95°C (15 s). The expression level was calculated using the delta threshold cycle value and is indicated as a value relative to the control.

### Measurement of [Ca^2+^]_i_

When the H1975 cells reached 80% confluence, they were transferred onto coverslips and incubated with 4 μM Fura2 and 0.05% pluronic acid in Reg (containing 147 mM Cl^—^, Table 1) for 15 min at RT in the dark. After incubation with Fura2, the cells were washed with Reg for 5 min before measuring [Ca^2+^]_i_. Solutions with different Cl^—^ concentrations were produced, as described in Table 1. The time course of the solutions applied is represented by bars above the traces. [Ca^2+^]_i_ was determined by measuring Fura2 fluorescence using dual excitation wavelengths of 340 nm and 380 nm and an emission wavelength of 530 nm. [Ca^2+^]_i_ is represented by the fura2 fluorescence ratio (340/380). The emitted fluorescence was monitored using a CCD camera (Retiga 6000, Q-imaging, Tuscon, AZ, USA) attached to an inverted microscope (Olympus, Japan) and analyzed using a Meta Fluor system (Molecular Devices, San Jose, CA, USA).

**Table 1.** Composition of solutions

| Composition (Reg) | Concentration (mM) |  |  |  |  |
| --- | --- | --- | --- | --- | --- |
| Sodium chloride (NaCl) | 140 |  |  |  |  |
| HEPES | 10 |  |  |  |  |
| Glucose | 10 |  |  |  |  |
| Potassium chloride (KCl) | 5 |  |  |  |  |
| Magnesium chloride (MgCl<sub>2</sub>) | 1 |  |  |  |  |
| Calcium chloride (CaCl<sub>2</sub>) | 1 |  |  |  |  |
| pH 7.4 |  |  |  |  |  |
| 310 mOsm (adjusted with mannitol) |  |  |  |  |  |
| Composition<br>(chloride solutions) | Concentration (mM) |  |  |  |  |
|  | 100 | 50 | 20 | 5 | 0 |
|  | Cl<sup>-</sup> | Cl<sup>-</sup> | Cl<sup>-</sup> | Cl<sup>-</sup> | Cl<sup>-</sup> |
| Sodium chloride (NaCl) | 100 | 50 | 20 | 5 | 0 |
| Sodium gluconate | 40 | 90 | 120 | 135 | 140 |
| HEPES | 10 |  |  |  |  |
| Glucose | 2 |  |  |  |  |
| Potassium gluconate | 5 |  |  |  |  |
| Magnesium sulfoxide (MgSO<sub>4</sub>) | 1 |  |  |  |  |
| Calcium gluconate | 1 |  |  |  |  |
| pH 7.4 |  |  |  |  |  |
| 310 mOsm (adjusted with mannitol) |  |  |  |  |  |

Fluorescence images, obtained at 3 s intervals, were normalized by subtracting the raw signals from the background images.

### Measurement of [Cl^—^]_i_

The H1975 cells were transferred onto coverslips after reaching 80% confluence and incubated with 5 mM MQAE and 0.05% pluronic acid in Reg for 15 min at RT in the dark. After incubation with MQAE dye, the cells were washed with Reg for 5 min prior to measuring [Cl^—^] _i_. The time course of applied solutions is represented with bars above the traces. [Cl^—^]_i_ was determined by measuring MQAE fluorescence using an excitation wavelength of 360 nm and an emission wavelength of 530 nm. The emitted fluorescence was collected with a CCD camera (Retiga 6000) attached to an inverted microscope (Olympus) and analyzed using a Meta Fluor system (Molecular Devices). Fluorescence images were obtained at 3 s intervals.

### Measurement of pH_i_

The pH_i_ of the H1975 cells was evaluated using BCECF fluorescence imaging. The cells were transferred onto coverslips after they had reached 80% confluence and incubated with 20 μM BCECF and the same volume of 0.05% pluronic acid to enhance loading efficacy in Reg for 15 min at RT in the dark. After incubation with BCECF dye, the cells were washed with Reg for 5 min prior to measuring pH_i_. The time course of the solutions applied is represented by the bars above the traces. pH_i_ was determined by measuring fluorescence using dual excitation wavelengths of 495 and 440 nm and an emission wavelength of 530 nm. pH_i_ is represented by the fluorescence ratio. Ratios of BCECF were converted to pH units using in situ calibration curves as described by Lee *et al*. (Lee *et al*, 2018). The calibration curve for pH_i_ is shown in Fig EV6A. Briefly, the BCECF-loaded cells were incubated in pH 5.5 calibration solution including 20 μM nigericin (Sigma; Cat: N7143) for 5 min, and then the process was repeated at pH 6.0-8.5 (0.5 interval). The pH value was calculated from the pH calibration curve using the following equation (pH = pKa-log((R_max_-R)/R-R_*min*_)) (pKa of BCECE; 6.97, R; ratio of BCECF, R_max_; maximum ratio, R_min_; minimum ratio). The emitted fluorescence was collected with a CCD camera (Retiga 6000) attached to an inverted microscope (Olympus) and analyzed using a Meta Fluor system (Molecular Devices). Fluorescence images were obtained at 3 s intervals.

### Measurement of cellular volume changes

When the confluence of H1975 cells reached 80%, they were transferred onto coverslips and incubated with 2 μM cell volume indicator calcein-AM and 0.05% pluronic acid in Reg for 15 min at RT in the dark. The calcein-loaded cells were washed with Reg for 5 min prior to measuring the volume changes. The time course of the solutions applied is represented by the bars above the traces. The cellular volume was determined by measuring the fluorescence at an excitation wavelength of 495 nm and an emission wavelength of 515 nm. The emitted fluorescence was collected with a CCD camera (Retiga 6000) attached to an inverted microscope (Olympus) and analyzed using a Meta Fluor system (Molecular Devices). Fluorescence images were obtained at 3 s intervals.

### Immunofluorescence and confocal microscopy

The transferred H1975 cells onto coverslips were incubated under the indicated conditions and then fixed with 4% paraformaldehyde (PFA) in PBS for 10 min at RT. The fixed cells were washed three times with PBS and then incubated with blocking serum (0.5% BSA and 5% goat serum in PBS) for 1 h at RT to block non-specific binding sites. The blocked cells were incubated with the primary antibodies overnight at 4°C, followed by three washes with PBS at RT. For the detection of the nuclear envelope, anti-Lamin A/C antibody (1:100 dilution in blocking serum) (Abcam, UK; Cat: ab185014) was used. Lysosomal vesicles were stained with anti-LAMP-2 mouse antibody (1:100) (Abcam; Cat: ab25631), which was detected by incubation with fluorescein isothiocyanate-tagged anti-mouse IgG antibody (green, 1:200) and rhodamine-tagged anti-mouse IgG antibody (red, 1:200) (Jackson ImmunoResearch, West Grove, PA, USA; Cat:115-095-003 and 115-025-072) for 1 h at RT. TFEB was stained with anti-TFEB rabbit antibody (1:100) (Cell Signaling Technology, Danvers, MA, USA; Cat:37785), which was detected by incubation with rhodamine-tagged anti-rabbit IgG antibody (red, 1:200) (Jackson ImmunoResearch; Cat:111-025-003) for 1 h at RT. After incubation, the cells were washed three times, and the cover slips were mounted on a glass slide with 20 μL of Fluoromount-G containing DAPI (Electron Microscopy Sciences, Hatfield, PA, USA; Cat:17984-24). Confocal microscopy images were obtained using an LSM 700 Zeiss confocal microscope (Carl Zeiss, Germany) with ZEN software (Carl Zeiss).

### Transmission electron microscopy

After incubation with Reg or 5Cl^—^ solution, H1975 cells were detached and fixed with 2% PFA and 2% glutaraldehyde in 100 mM phosphate buffer (pH 7.4) for 12 h at 4°C. Fixed cells were washed with 100 mM phosphate buffer and post-fixed with 1% osmium tetroxide in 100 mM phosphate buffer for 2 h at RT. Thereafter, the cells were dehydrated with gradually increasing concentrations of ethanol (50–100% in water). The cells were infiltrated with propylene oxide for 10 min and embedded using a Poly/Bed 812 kit (Polysciences, Warrington, PA, USA; Cat:08792). The specimens were sectioned at 200 nm using an ultramicrotome (EM-UCT, Leica, Teaneck, NJ, USA) and stained with toluidine blue. The specimens were sectioned at 70 nm and stained with 5% uranyl acetate for 15 min, and then with 1% lead citrate for 7 min. TEM images were obtained using a transmission electron microscope (JEM-1011, JEOL, Japan) at an acceleration voltage of 80 kV.

### Western blotting

The cells incubated under the indicated conditions were dispersed in lysis buffer (Cell Signaling; 9803) containing 20 mM Tris, 150 mM NaCl, 2 mM EDTA, 1% Triton X-100, and protease inhibitor mixture in the presence (for phosphorylated form) or absence of phosphatase inhibitor mixture. The cell lysate was sonicated and centrifuged at 11,000 × g for 15 min at 4°C. Protein concentration was calculated using the Bradford assay kit (Bio-Rad, Hercules, CA, USA; Cat:5000001). The obtained protein samples were incubated with sodium dodecyl sulfate (SDS) protein sample buffer, and then the samples were separated by SDS-polyacrylamide gel electrophoresis, followed by transfer onto polyvinylidene difluoride membranes soaked in methanol. The membranes were incubated for 1 h at RT with 5% non-fat milk solution in 0.05% Tris-buffered saline with 0.05% Tween-20 to block non-specific binding. The blocked membranes were incubated overnight with primary antibodies at 4°C. CLC7 was detected using anti-CLC7 rabbit antibodies (Novusbio, Centennial, CO, USA; Cat: NBP2-30021). TRPML1 was detected using anti-TRPML1 (MCOLN1) rabbit antibody (ATLAS antibodies, Sweden; Cat: HPA031763). Cleaved PARP was detected using an anti-cleaved PARP (Asp214) antibody (Cell Signaling; Cat:9541). mTORC1s, including both total and phosphorylated forms, were detected using anti-mTOR, anti-phospho-mTOR (Ser 2448) rabbit antibody and anti-phospho-mTOR (Ser 2481) rabbit antibodies (Cell Signaling; Cat:2972, 2971, and 2974). TPC1 and TPC2 were detected using anti-TPCN1 and anti-TPCN2 rabbit antibodies (Alomone Labs; Cat: ACC-071 and ACC-072). After primary antibody incubation, the membranes were washed with PBS three times and then incubated with secondary antibodies: horseradish peroxidase-conjugated anti-mouse IgG and anti-rabbit IgG antibodies (Millipore, Billerica, MA, USA; Cat: AP124P and AP132P) for 1 h at RT. β-actin was detected using a horseradish peroxidase-conjugated anti-β-actin mouse antibody (Sigma; Cat: A3854). The membrane was washed three times, and the protein was detected with enhanced chemiluminescence on X-ray films.

### Transwell membrane migration assay

The dispersed H1975 cells (5 × 10^4^ cells/well) in Reg, 5 mM Cl^—^ solutions, or RPMI (with 1% FBS each) were added to the upper chamber of a 6-well transwell membrane plate (8.0 μm pore sized insert). The bottom chambers were filled with pH 7.5 or pH 6.5 RPMI (with 10% FBS and 100 U/mL penicillin-streptomycin), followed by the indicated conditions. After incubation for 3 h, the membranes were stained with DAPI (blue) or crystal violet (purple). The membranes were incubated with chilled methanol (preserved at -20°C) for 1 min at -20°C to fix the cells, and then washed with PBS three times. For DAPI staining, the membranes were incubated with DAPI solution (1μg/mL in distilled water (DW)) for 30 min at 37°C in the dark and then washed twice with DW. For crystal violet staining, the membranes were incubated with 0.25% crystal violet solution in DW (with 20% methanol) for 10 min at room temperature in the dark, and then washed with DW six times. After washing the membranes, the media on the top was carefully removed, and DW was added to the bottom of the plate. The plates were subsequently analyzed using an LSM 700 Zeiss confocal microscope (Carl Zeiss) (for DAPI) or an inverted microscope (Olympus) with Mosaic software (Opto Science, Japan) (for crystal violet). The intensity of the obtained images was measured using the Meta Morph system (Molecular Devices).

### Flow cytometry

To analyze cell viability, H1975 cells were treated with Reg and 5 mM Cl^—^ solution under the indicated conditions (0.5, 1, 6, 12, and 24 h). Thereafter, the cells were washed with annexin V binding buffer (50 mM HEPES, 700 mM NaCl, 12.5 mM CaCl_2_, pH 7.4) and suspended in 100 μL of annexin V binding buffer. A single-cell suspension was treated with Pacific Blue-conjugated annexin V (Thermo; Cat: A35122) and propidium iodide (PI, Thermo; Cat: P1304MP) for 15 min at room temperature in the dark. The negative control was incubated with annexin V-binding buffer without annexin V and PI. After incubation, 500 μL of annexin V buffer was added to the samples, and the samples were analyzed at 410 nm excitation with a 455 nm band-pass filter to detect Pacific blue and 535 nm excitation with a 620 nm band-pass filter to detect PI. The percentage of late apoptosis was calculated by analyzing the first quadrant of the cell population, which was determined using annexin V and PI fluorescence.

### pHRodo staining

The transferred H1975 cells onto coverslips were incubated with Reg and 5 mM Cl^—^ solution for 30 min at 37°C and then fixed with 4% PFA in PBS for 10 min at RT. The fixed cells were washed with PBS three times, and then treated with a pHRodo Green AM Intracellular pH Indicator (Invitrogen; Cat: P35373) in Reg for 30 min at 37°C according to the manufacturer’s instructions. After incubation, for the measurement of pH calibration, the cells were incubated with different pH values of Reg (4.5, 5.5, 6.5, and 7.5). The calibration curve of pHRodo is shown in Fig EV6B. The coverslips were mounted on glass slides, and images were obtained using an LSM 700 Zeiss confocal microscope (Carl Zeiss, Germany) with ZEN software (Carl Zeiss).

### Statistical analyses

Data from the indicated number of experiments were expressed as mean ± standard error of the mean (SEM). Statistical significance was determined by analysis of variance for each experiment (*p < 0.05, **p < 0.01, *** p < 0.001), which was analyzed using ANOVA (followed by a Newman-Keuls multiple comparison test).

## Abbreviations

340/380: fura2 fluorescence ratio
Baf: bafilomycin A1
BAPTA: 1,2-bis(o-aminophenoxy)ethane-N,N,N′,N′-tetraacetic acid
[Ca^2+^]_i_: intracellular Ca^2+^ concentration
CBM: chloride binding motif
[Cl^—^]_i_: intracellular Cl^—^ concentration
CPA: cyclopiazonic acid
DAPI: 4’,6-diamidino-2-phenylindole
DW: distilled water
EGFR: epidermal growth factor receptor
FBS: fetal bovine serum
Fura: Fura-2-AM
GPN: glycyl-Phenylalanine β-naphthylamide
MQAE: ethoxycarbonylmethyl-6-methoxyquinolinium bromide
NPPB: 5-Nitro-2-(3-phenylpropylamino) benzoic acid
PCR: polymerase chain reaction
PFA: paraformaldehyde
pHi: intracellular pH
PI: propidium iodide
Reg: physiological salt solution
RPMI: Roswell Park Memorial Institute 1640
RT: room temperature
SDS: sodium dodecyl sulfate
siRNA: small interfering RNA
TEM: transmission electron microscopy
TFEB: transcription factor EB
TPC: two pore channel.

## Acknowledgements

This work was supported by a National Research Foundation of Korea (NRF) grant funded by the Korean government [MSIT; NRF-2022R1A2C1003890 (JHH) and NRF-2020R1A2C2003409 (DMS)]. All image data acquisition was performed at the Cell to In Vivo Imaging Core Facility Research Center (CII, Lee Gil Ya Cancer and Diabetes Institute, Gachon University, Incheon, South Korea).

## Author contributions

J.H.H, D.M.S, and D.L. conceptualized and designed the study and acquired, analyzed, and interpreted the data. D.L. and J.H.H. drafted the manuscript and acquired the data. D.M.S. revised the manuscript critically for important intellectual content. D.L. and J.H.H. drew all schematic illustrations. J.H.H. and D.M.S. contributed to the final approval of the version to be published and agreed to be accountable for all aspects of the work in ensuring that questions related to the accuracy or integrity of any part of the work are appropriately investigated and resolved.

## Figure legends

**Expanded View Figure 1.**
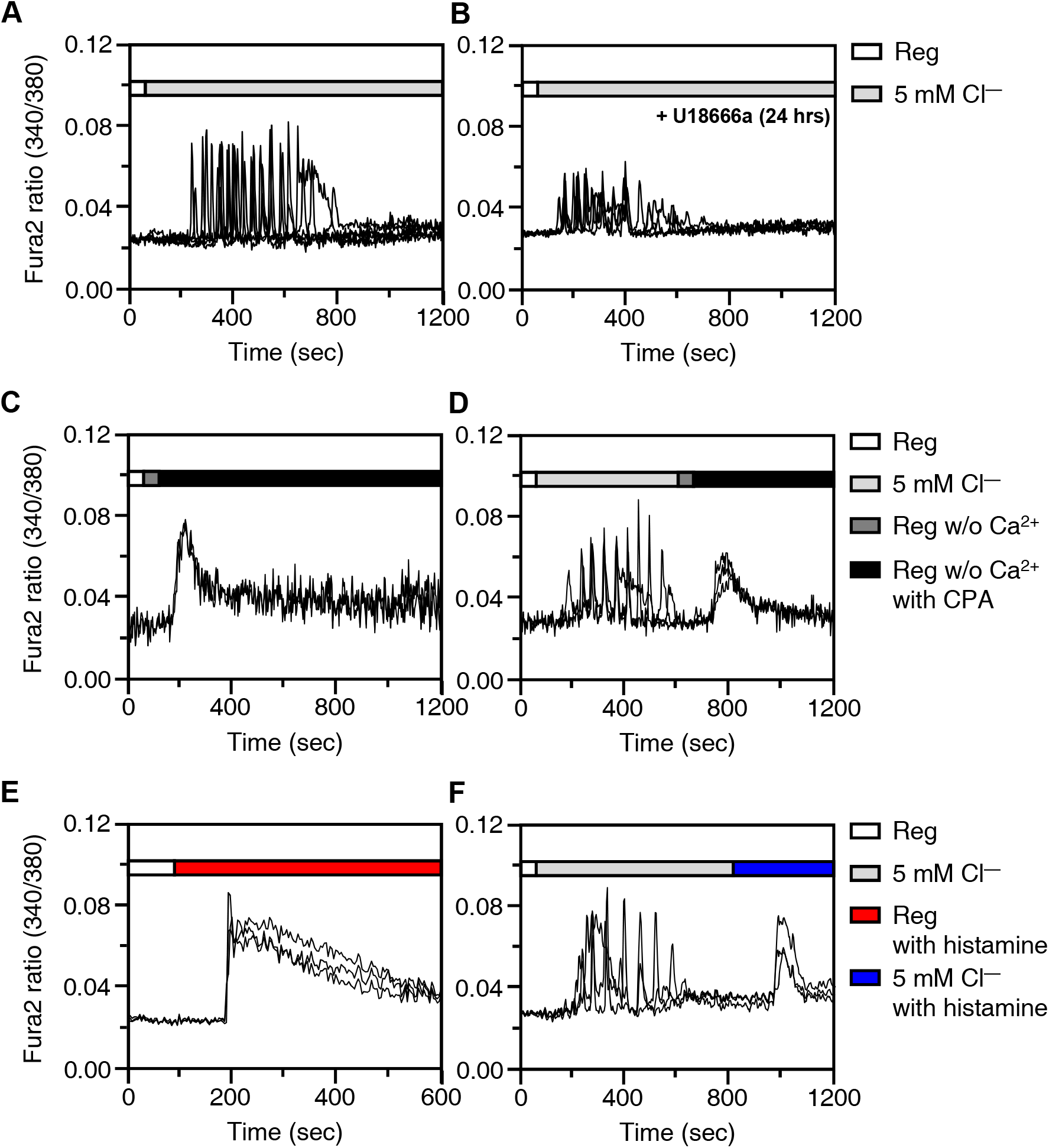
The effect of low Cl^—^-mediated Ca^2+^ signaling on ER Ca^2+^ release. **(A, B)** Changes in [Ca^2+^]_i_ in H1975 cells with 5 mM Cl^—^-induced Ca^2+^ signaling in the absence of U18666a (4 μg/mL, 24 hrs) or in the presence of U18666a. **(C, D)** Changes in [Ca^2+^]_i_ in H1975 cells in CPA treatment (A) and 5 mM Cl^—^-induced Ca^2+^ signaling with subsequent treatment of CPA (B). **(E, F)** Changes in [Ca^2+^]_i_ in H1975 cells in histamine treatment (C) and 5 mM Cl^—^-induced Ca^2+^ signaling with subsequent treatment of histamine (D).

**Expanded View Figure 2.**
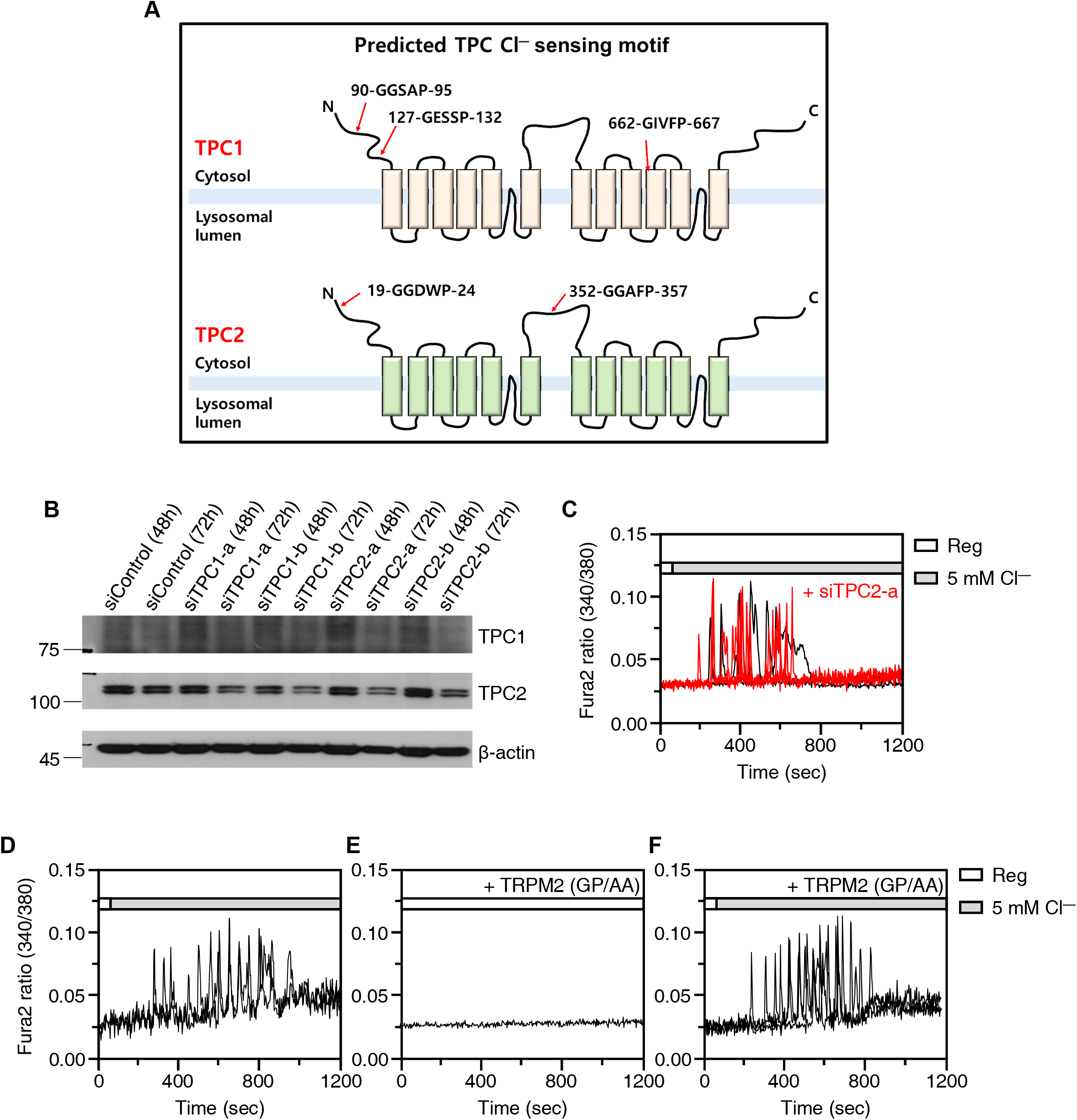
Modulation of TPC expression and TRPM2 GXXXP motif have no effect on low Cl^—^-induce Ca^2+^ signaling. **(A)** Schematic illustration of TPCs structure marked with GXXXP motif. **(B)** Western blotting analysis of TPC1 and TPC2 in siTPC1- or siTPC2-transfected H1975 cells. **(C)** Changes in [Ca^2+^]_i_ in siTPC2-a-transfected H1975 cells with 5 mM Cl^—^ solution. **(D-F)** Changes in [Ca^2+^]_i_ in H1975 cells with 5 mM Cl^—^ solution (A), Reg with TRPM2-GP/AA (B), 5 mM Cl^—^ with TRPM2-GP/AA (C).

**Expanded View Figure 3.**
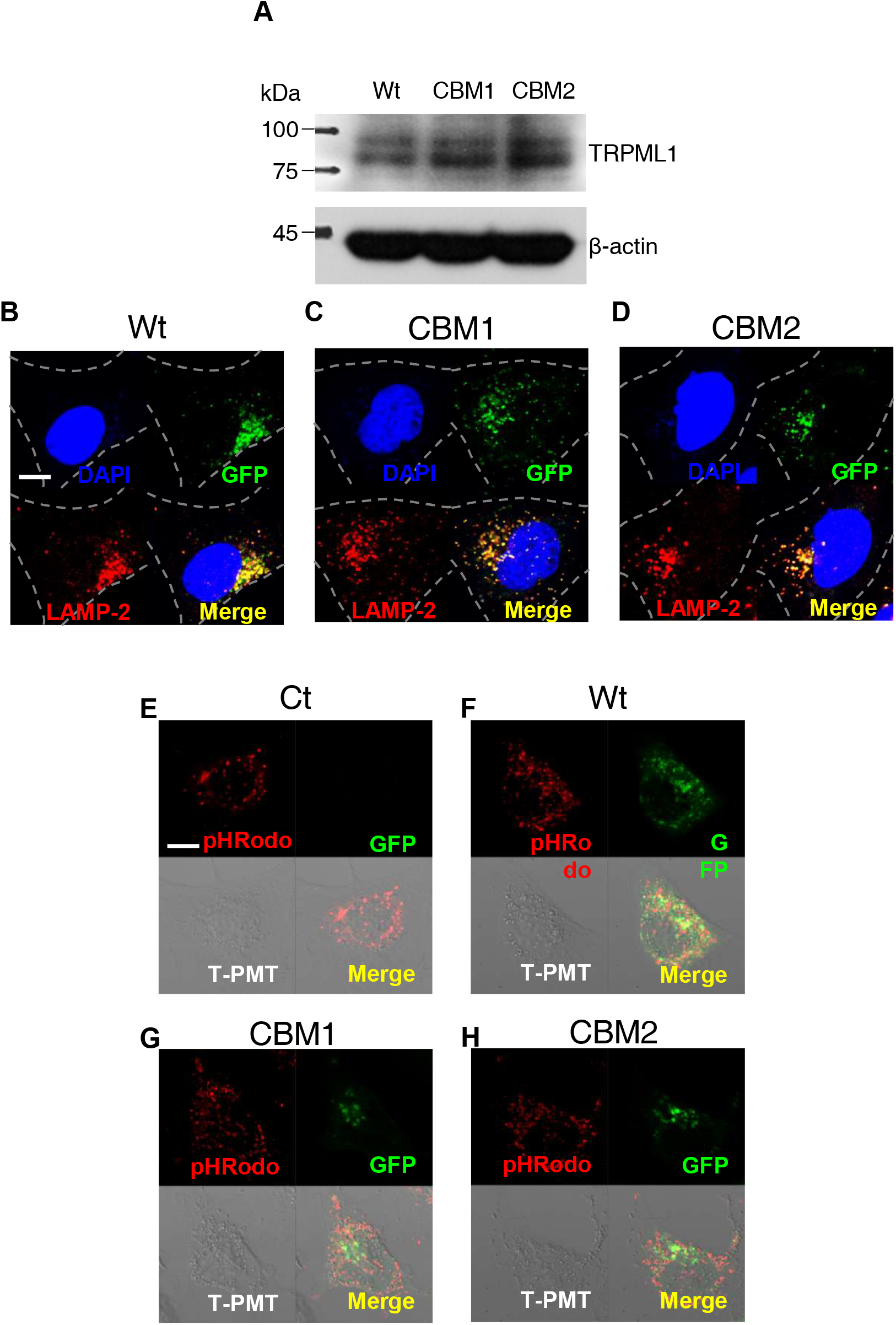
The effect of TRPML1 clones on LAMP2 location and vesicle pH. Western blotting analysis of TRPML1 protein in H1975 cells transfected with TRPML1 Wt, CBM1, and CBM2. **(B)** Confocal images of immunofluorescence staining of LAMP-2 (red) and DAPI (blue) which transfected with GFP-tagged TRPML1 Wt, CBM1, and CBM2 (green). The scale (white) represents 10 μm. **(E-H)** Confocal images of immunofluorescence staining of pHRodo (red) in control (E) and cells transfected with GFP-tagged TRPML1 Wt (F), CBM1 (G), or CBM2 (H). The scale (white) is 10 μm.

**Expanded View Figure 4.**
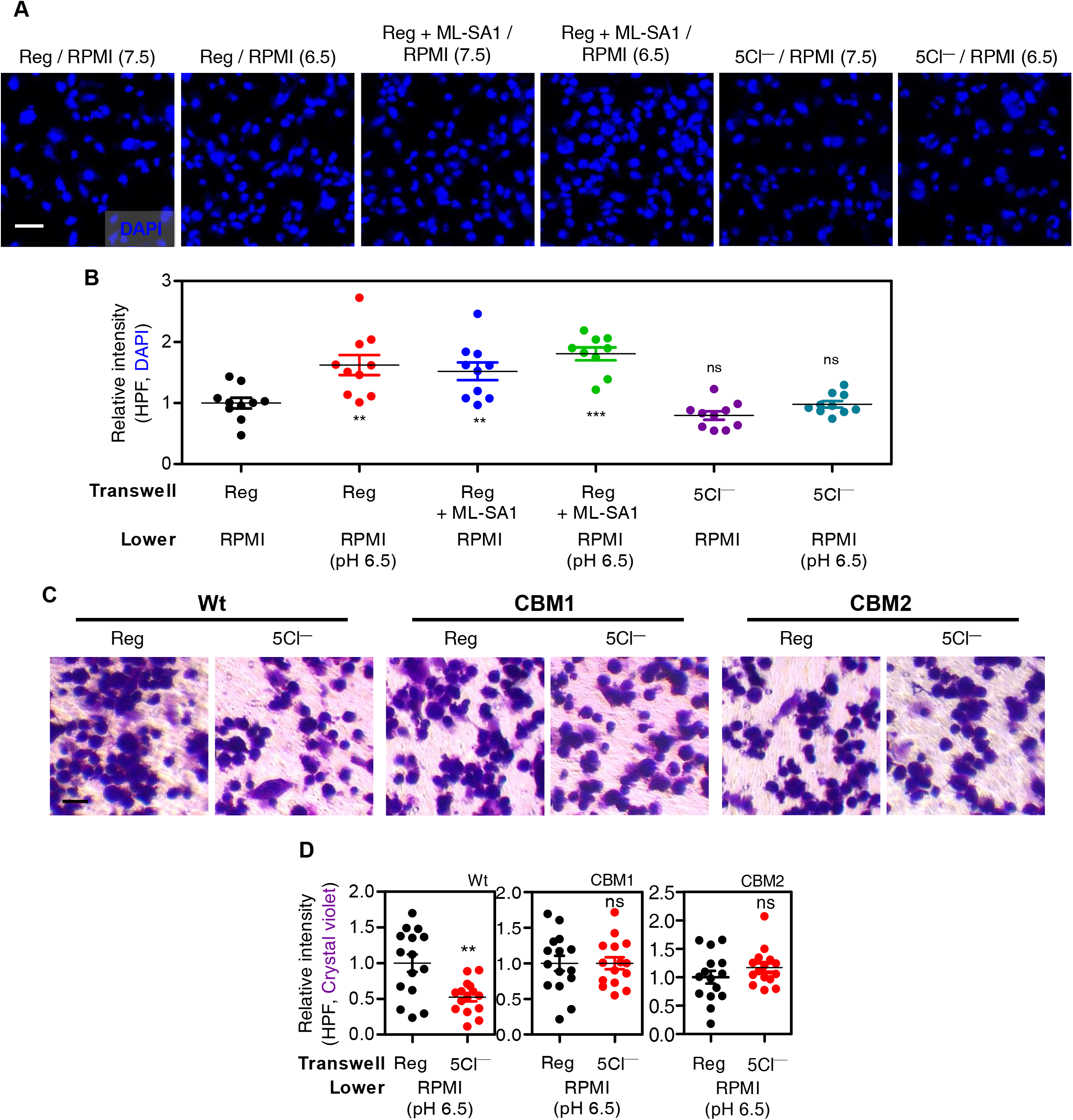
Transwell migration assay in presence of ML-SA1 or TRPML1 clones. **(A)** Images of transwell migration assay with DAPI (blue) or crystal violet (purple) which incubated for under the indicated conditions. **(B)** The dot plots are presented as means ± SEMs of the relative intensity of DAPI and crystal violet (**p < 0.01, ns; non-significance). **(C)** Images of transwell migration assay with crystal violet (purple) in TRPML1, CBM1, and CBM2-transfected cells with the indicated conditions. **(D)** The dot plots are presented as means ± SEMs of the relative intensity of DAPI and crystal violet (*p < 0.05).

**Expanded View Figure 5.**
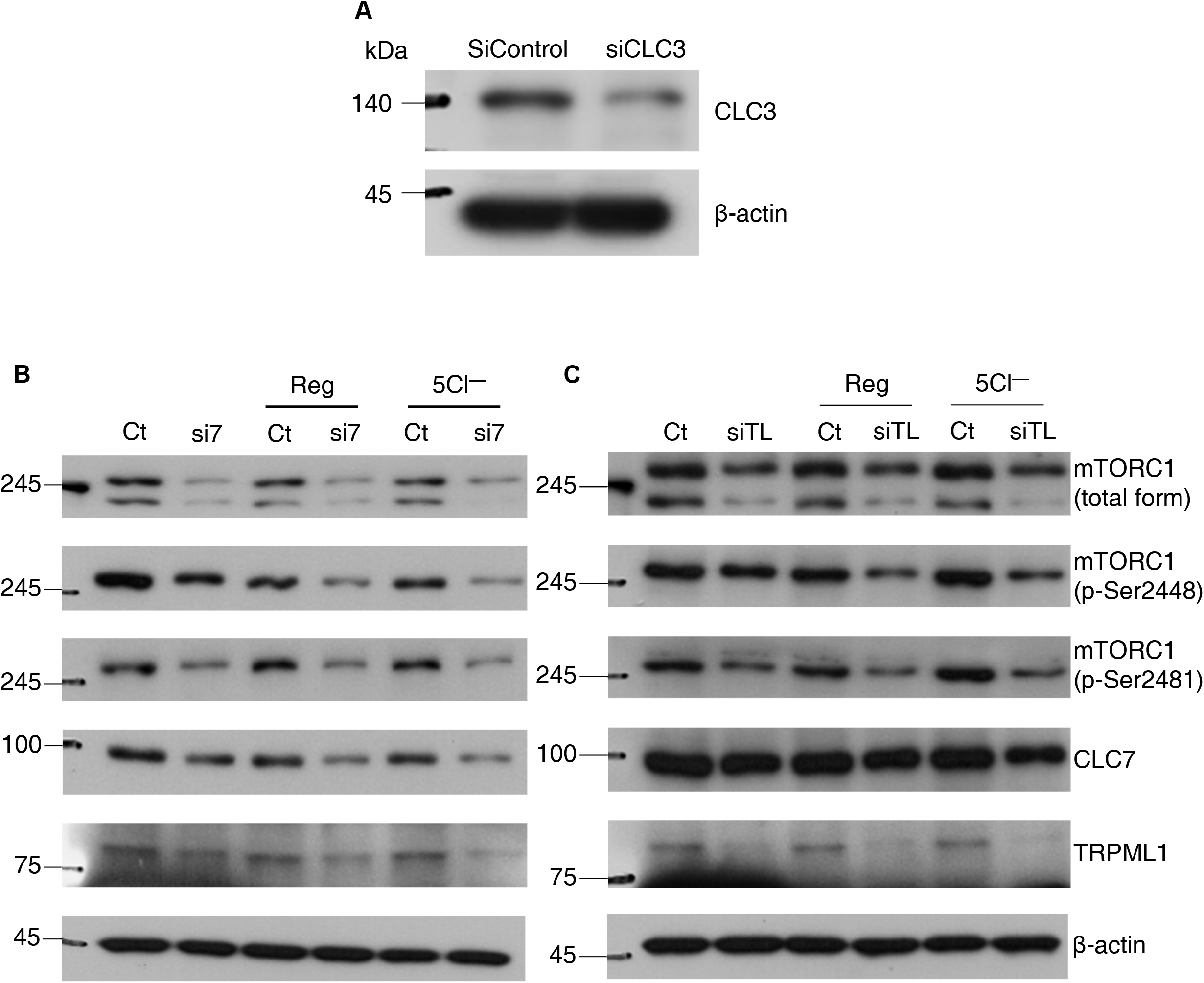
Western blotting analysis with siCLC3, siCLC7 and siTRPML1. **(A)** Western blotting analysis of CLC3 transfected with siRNA-CLC3 (siCLC3). **(B)** Western blotting analysis of mTORC1, phosphorylated mTORC1 (ser-2448 and ser-2481), CLC7 and TRPML1 transfected with siRNA-CLC7 (si7) in presence of Reg or 5 mM Cl^—^ solution at indicated time (30 min). **(C)** Western blotting analysis of mTORC1, phosphorylated mTORC1 (ser-2448 and ser-2481), CLC7 and TRPML1 transfected with siRNA-TRPML1 (siTL) in presence of Reg or 5 mM Cl^—^ solution at indicated time (30 min).

**Expanded View Figure 6.**
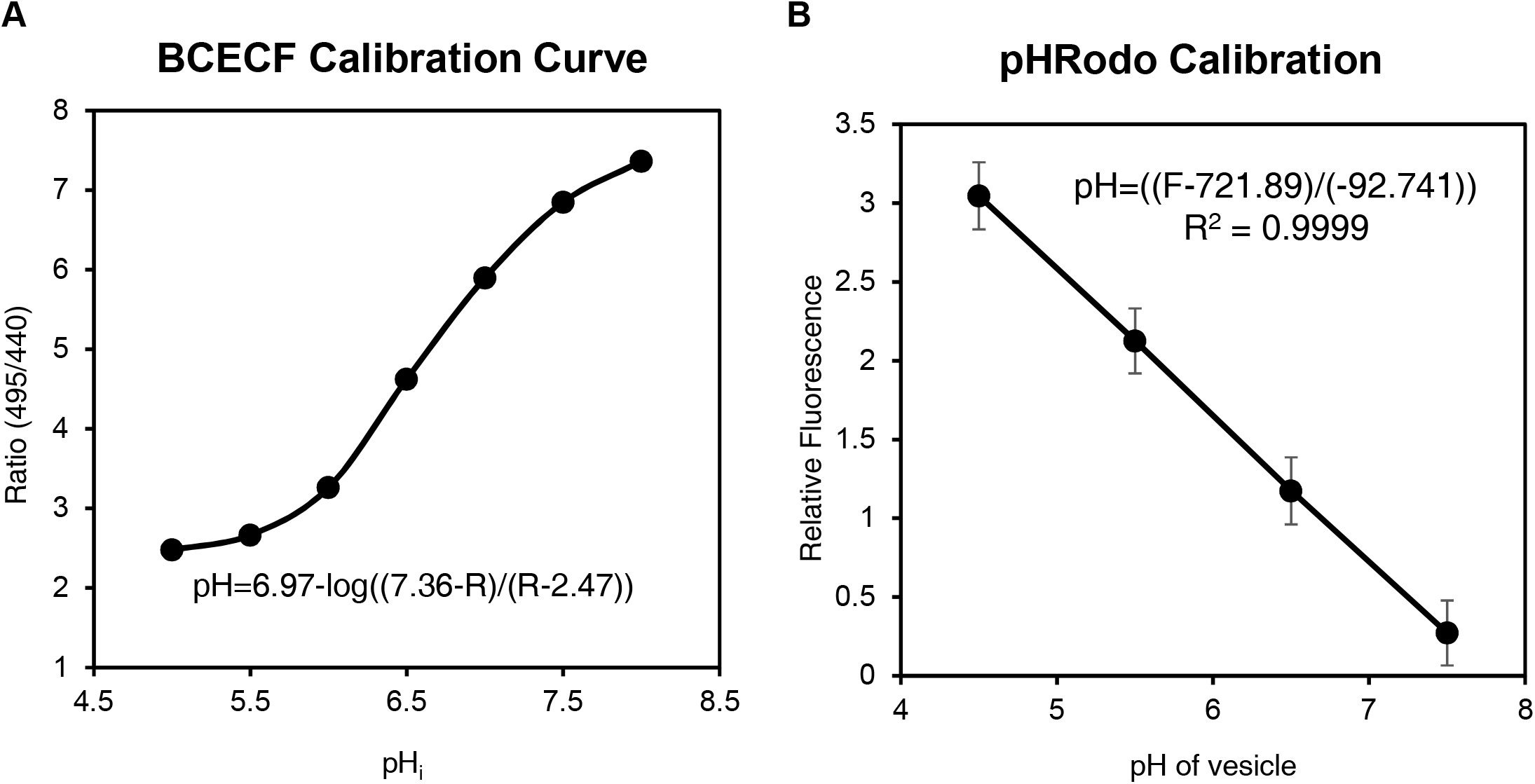
pH calibration of BCECF and pHRodo. **(A)** The pH calibration curves for BCECF-loaded H1975 cells at pH 5.5, 6.0, 6.5, 7.0, 7.5, 8.0, and 8.5. The regression curve was pH=6.97-log((7.36-R)/(R-2.47)), where R is the ratio of BCECF fluorescence. **(B)** The pH calibration curve for pHRodo-loaded H1975 cells at pH 4.5, 5.5, 6.5, and 7.5. Regression curves are pH =((F-721.89)/(−92.741)), F: intensity of pHRodo fluorescence.

